# Allopeptimicins: unique antibacterial metabolites generated by hybrid PKS-NRPS, with original self-defense mechanism in *Actinoallomurus*

**DOI:** 10.1101/2022.04.01.486743

**Authors:** Marianna Iorio, Andrea Gentile, Cristina Brunati, Arianna Tocchetti, Paolo Landini, Sonia Ilaria Maffioli, Stefano Donadio, Margherita Sosio

## Abstract

In the search for structurally novel metabolites with antibacterial activity, innovative approaches must be implemented to increase the probability of discovering novel chemistry from microbial sources. Here we report on the application of metabolomic tools to the genus *Actinoallomurus*, a poorly explored member of the Actinobacteria. From examining extracts derived from 88 isolates belonging to this genus, we identified a family of cyclodepsipeptides acylated with a C_20_ polyketide chain, which we named allopeptimicins. These molecules possess unusual structural features, including several double bonds in the amino-polyketide chain and four non-proteinogenic amino acids in the octapeptide. Remarkably, allopeptimicins are produced as a complex of active and inactive congeners, the latter carrying a sulfate group on the polyketide amine. This modification is also a mechanism of self-protection in the producer strain. The structural uniqueness of allopeptimicins is reflected in a biosynthetic gene cluster showing a mosaic structure, with dedicated gene cassettes devoted to formation of specialized precursors and modular assembly lines related to those from different pathways.

## Introduction

The spread of antibiotic resistance is a serious threat to public health that is forecast to reach dramatic levels by 2050, exacerbated by the scarcity of new antibacterial drugs in the development pipeline.^1^ Most of the antibiotics in clinical use today are microbial metabolites or derivatives thereof,^2^ while alternative approaches to antibiotic discovery have so far failed to be as successful as microbial product screening.^2^ However, through several decades of large screening efforts, thousands of metabolites with antibacterial activity have been reported in the literature, rendering the discovery of new scaffolds by bioactivity-based screening a labor-intensive endeavor.

It is generally believed that the known metabolites represent the tip of the iceberg of molecules produced by microorganisms and that alternative approaches can reduce the biases imposed by bioassay-based screens.^1,3,4^ An attractive approach is represented by employing mass-spectrometry (MS)-based metabolomic tools to navigate the chemical complexity observed in microbial fermentation extracts. Recent advancements in this area are represented by molecular networking tools^5^ and by machine learning that, combined with publicly accessible databases, have greatly expedited metabolite annotation and prioritization for further investigations.^6,7^ Particularly promising is the combination of metabolomic tools to poorly explored bacterial taxa. In recent work on bacterial strains belonging to the actinobacterial genus *Planomonospora*, we uncovered novel chemistry, which led to a family of unexpected biosynthetic gene clusters (BGCs),^8,9^ which in turn helped uncover further novel chemistry.^10^

We previously reported that strains belonging to another actinobacterial genus, *Actinoallomurus*, produce at high frequency molecules with antibacterial activity,^11^ with several new variants of known scaffolds identified through bioactivity-based screening.^12,13,14,15,16^ Here we report that metabolomic analysis of *Actinoallomurus* spp. unveiled an unprecedented acylated cyclodepsipetide with unusual features and potent antibacterial activity.

## Results and Discussion

### Discovery of allopeptimicins

Microbial fermentation extracts derived from 88 *Actinoallomurus* strains from the NAICONS collection were analyzed by low resolution LC-MS/MS followed by molecular networking (Fig. S1), which evidenced previously reported metabolites such as the polyether α-823,^11^ the paramagnetoquinones,^12^ the allocyclinones,^13^ and the isoflavonoids genistein and daidzein.^16^

Among all the observed clusters, one molecular family attracted our attention: it was associated with an extract generated from strain ID145808 only and was absent in extracts from the remaining 87 strains. This molecular family consisted of a group of metabolites characterized by a parent mass of around 1200 amu (Fig. S1), whose fragmentation spectra showed no matches to the 11,600 MassIVe datasets present in the GNPS repository (https://gnps.ucsd.edu/) and to our private spectral library of metabolic profiles from about 14,000 extracts. Interestingly, the microbial extract coming from strain ID145808 possessed activity against *Staphylococcus aureus*.

HPLC fractionation of the ID145808 extract followed by bioassay of the resulting fractions enabled signal deconvolution: a group of related molecules with *m/z* of 1205, 1207, 1219 and 1221 [M+H]^+^ and elution times of 6.76, 6.79, 6.90 and 7.07 min, respectively (Fig. 1). Thus, the four related molecules could be accounted for by mass differences of 2 and 14 amu. Inspection of the LC-MS profile of the entire extract revealed a second group of four related molecules with mass distances of 2 and 14 amu. This second group of molecules, eluting between 7.53 and 8.05 min, was present in HPLC fractions devoid of antibacterial activity (Fig. 1) and showed *m/z* of 1285, 1287, 1299 and 1301 [M+H]^+^. Thus, molecules from one group differed by 80 amu from molecules of the other group. All molecules from both groups shared the same UV spectrum with four maxima: the first at 239 nm (ε/dm^3^ mol^-1^ cm^-1^ 23700) and a distinctive tricuspid absorption at 260 (25100), 270 (31000) and 280 nm (24200), suggestive of three conjugated double bonds. High Resolution mass spectrometry (HR-MS) analysis resolved the distances among the congeners as two hydrogens, a methylene group and a sulphate moiety, as indicated by the molecular formulas reported in Table 1. The active and inactive congeners were named allopeptimicin A1 through A4 and B1 through B4, respectively. The most abundant congener in each group were A3 and B4.

**Table 1.**
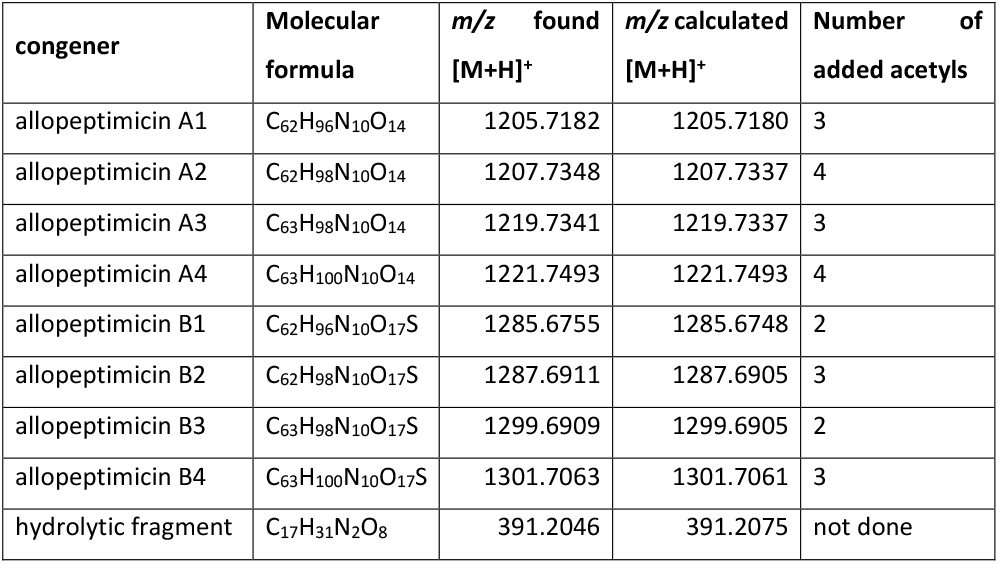
Properties of allopeptimicin congeners and fragments thereof, and number of added acetyls

**Fig. 1.**
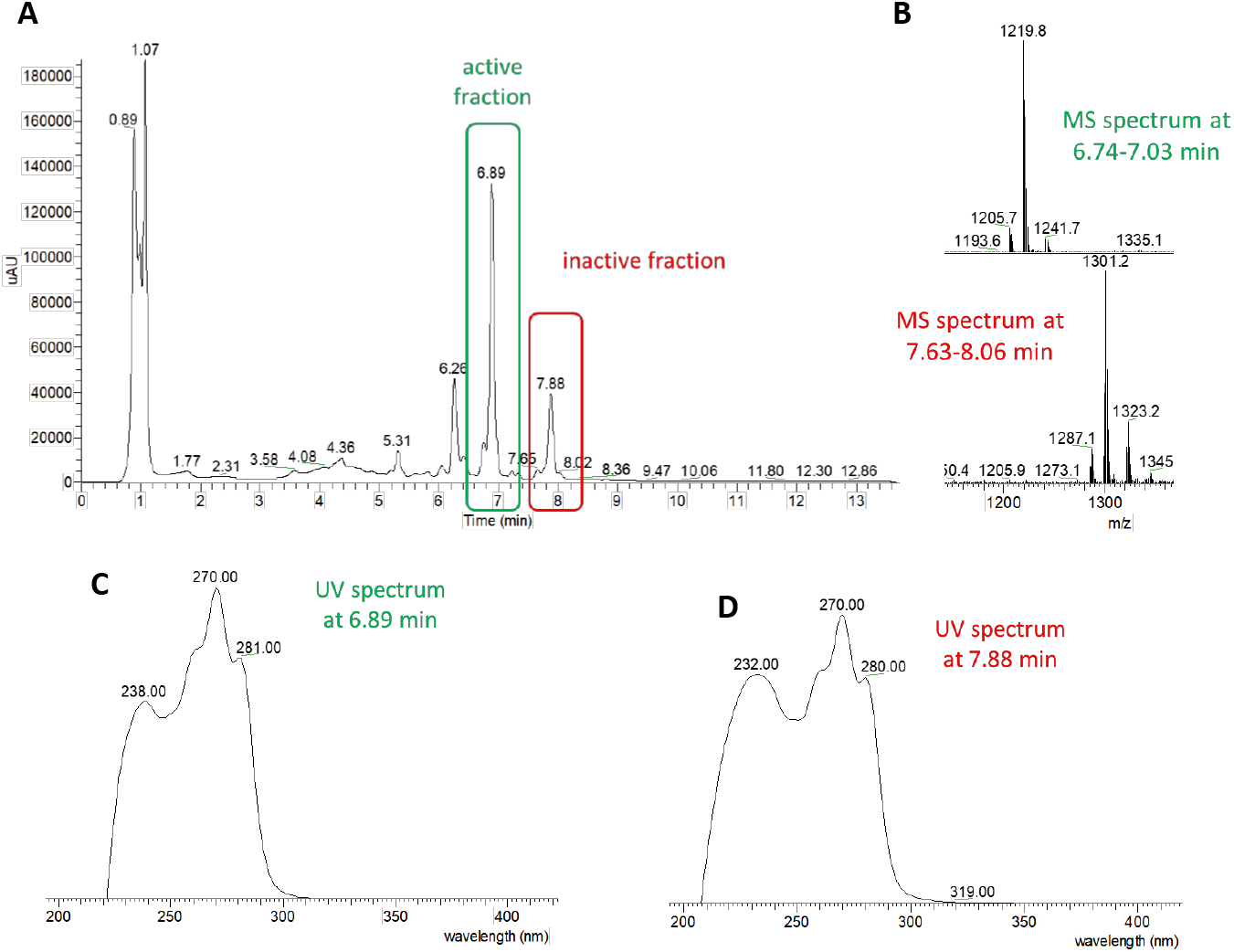
A) HPLC profile of the extracted fermentation broth from a culture of *Actinoallomurus* ID145808. The green and red boxes denote the regions of the chromatogram containing fractions with and devoid of antibacterial activity, respectively. B) MS of the highlighted regions in A). C-D) UV−vis spectra of the fractions at 6.89 and 7.88 minutes, respectively.

### Structure Elucidation

The molecular formulas together with the peculiar UV absorption of allopeptimicins suggested a peptide-polyketide skeleton: the peptide portion would be consistent with the presence of several nitrogen and oxygen atoms, while the large number of carbons and the conjugated double bonds predicted from the UV spectrum pointed to a long hydrocarbon portion. Under alkaline conditions all allopeptimicin congeners underwent hydrolysis with the elimination of 390 amu after the addition of two water molecules, suggesting the presence of two hydrolysable bonds, likely esters, in a cyclic structure (Fig. S2A and S2B). Amidation with methylamine led to the formation of one methylamide in each congener (Fig. S2C and S2D), revealing the presence of one free carboxylic acid. Acetylation reactions, carried out in order to establish the number of free amines or hydroxyls, showed differences among congeners, with the incorporation of two, three or four acetyls, with a clear pattern (Table 1): congener pairs differing for one methylene group (A1 and A3, B1 and B3, A2 and A4, and B2 and B4) incorporated an identical number of acetyls while, in congeners differing by 2 amu, the smaller congener always incorporated one lesser acetyl group, indicating that the unsaturation in A1, B1, A3 and B3 involves an amine or a hydroxyl. Members of the B group incorporated one lesser acetyl than the corresponding members of the A group, suggesting that the sulphate moiety is present on an amine. [A sulphate ester is expected to hydrolyze under alkaline conditions.]

These properties of allopeptimicins guided interpretation of extensive 1D- and 2D-NMR experiments and HR-tandem MS analyses. A set of NMR experiments (Table 2 and Fig.s S3-S7) in CD_3_CN/D_2_O 8:2 was performed on a sample containing the A congeners in the following proportion: A1:A2:A3:A4 15:10:45:30.

**Table 2.** ^1^H and ^13^C NMR Spectral Data of Allopeptimicin A3 and B3 Measured at 300 MHz in CD_3_CN:D_2_O:H_2_O 4:1:0.2.

|  |  | Allopeptimicin A3 |  |  | Allopeptimicin B3 |  |
| --- | --- | --- | --- | --- | --- | --- |
| unit | position | δH (multiplicity, J coupling when visible (Hz)) | δC | NOE correlations | δH (multiplicity, J coupling when visible (Hz) ) | δC |
| Ile_1 | NH | 7.01 |  | 1.76 (methyl in C2-acyl chain) | 6.97 |  |
|  | 1 |  | 172.3 |  |  | 172.3 |
|  | 2 | 4.34 (t, 7.0) | 57.4 |  | 4.34 (t, 7.0) | 57.4 |
|  | 3 | 1.86 | 36.7 |  | 1.86 | 36.7 |
|  | 4a | 1.32 | 25.9 |  | 1.33 | 25.9 |
|  | 4b | 1.10 |  |  | 1.08 |  |
|  | 5 | 0.85 (d, 7.0) | 11.0 |  | 0.86 (d, 7.0) | 11.0 |
|  | 6 | 0.84 | 14.1 |  | 0.84 | 14.1 |
| Val_2 | NH | 7.58 |  | 4.34 (C2-Ile_1) | 7.56 |  |
|  | 1 |  | 171.4 |  |  | 171.4 |
|  | 2 | 4.16 | 58.9 |  | 4.16 | 58.9 |
|  | 3 | 2.00 | 30.8 |  | 2.00 | 30.6 |
|  | 4 | 0.86 (d, 7.0) | 17.6 |  | 0.85 (d, 7.0) | 18.8 |
|  | 5 | 0.87 (d, 7.0) | 18.6 |  | 0.90 (d, 7.0) | 18.6 |
| Thr_3 | NH | 7.76 |  | 4.16 (C2-Val_2) | 7.80 |  |
|  | 1 |  | 169.7 |  |  | 169.7 |
|  | 2 | 4.81 | 57.1 |  | 4.86 | 56.7 |
|  | 3 | 5.33 | 70.9 |  | 5.30 | 70.8 |
|  | 4 | 1.22 | 17.7 |  | 1.20 | 17.3 |
| Piz_4 | 1 |  | 172.4 |  |  | 172.4 |
|  | 2 | 5.46 | 50.3 |  | 5.46 | 50.7 |
|  | 3a | 2.09 | 25.4 |  | 2.10 | 25.4 |
|  | 3b | 1.84 |  |  | 1.78 |  |
|  | 4a | 1.53 | 20.7 |  | 1.53 | 20.7 |
|  | 4b | 1.49 |  |  | 1.46 |  |
|  | 5a | 2.99 | 46.3 |  | 3.00 | 46.1 |
|  | 5b | 2.69 |  |  | 2.73 |  |
| Piz_5 | 1 |  | 170.8 |  |  | 170.8 |
|  | 2 | 5.15 | 50.6 |  | 5.17 | 50.7 |
|  | 3a | 2.31 | 18.1 |  | 2.31 | 18.1 |
|  | 3b | 1.99 |  |  | 1.97 |  |
|  | 4a | 2.25 | 20.1 |  | 2.22 | 20.2 |
|  | 4b | 1.98 |  |  | 2.07 |  |
|  | 5 | 6.96 | 145.6 |  | 6.98 | 145.5 |
| Hmg_6 | 1 |  | 169.7 |  |  | 169.7 |
|  | 2 | 4.99 | 73.3 |  | 5.09 | 72.9 |
|  | 3a | 2.01 | 34.1 |  | 2.00 | 34.2 |
|  | 3b | 1.99 |  |  | 1.99 |  |
|  | 4 | 2.59 | 35.3 |  | 2.60 | 35.2 |
|  | 5 |  | 178.0 |  |  | 178.0 |
|  | 4-Me | 1.15 (d, 6.8) | 16.8 |  | 1.15 (d, 6.8) | 16.8 |
| Thr_7 | NH | 7.73 |  | 4.99 (C2-Hmg_6) | 7.91 |  |
|  | 1 |  | 171.6 |  |  | 171.6 |
|  | 2 | 4.70 | 54.5 |  | 4.72 | 54.5 |
|  | 3 | 3.89 | 67.5 |  | 3.88 | 67.7 |
|  | 4 | 1.09 (d, 6.0) | 18.9 |  | 1.13 (d, 6.0) | 19.1 |
| N-Me-Leu_8 | 1 |  | 171.5 |  |  | 171.5 |
|  | 2 | 5.15 | 53.2 |  | 5.16 | 53.2 |
|  | 3a | 1.69 | 37.2 | 3.02 (N-Me Leu_8) | 1.70 (m) | 37.3 |
|  | 3b | 1.52 |  |  | 1.54 (m) |  |
|  | 4 | 1.45 | 24.3 |  | 1.46 (m) | 24.2 |
|  | 5 | 0.81 (d, 6.8) | 20.6 |  | 0.81 (d, 6.8) | 20.5 |
|  | 6 | 0.87 (d, 6.8) | 22.7 |  | 0.89 (d, 6.8) | 18.8 |
|  | N-Me | 3.02 (s) | 30.8 |  | 3.02 (s) | 30.7 |
| Acyl chain | 1 |  | 170.4 |  |  | 170.4 |
|  | 2 |  | 130.9 |  |  | 130.9 |
|  | 3 | 6.25 (d, 9.0) | 136.2 |  | 6.25 (d, 9.0) | 136.2 |
|  | 4 | 2.21 | 27.7 |  | 2.21 | 27.7 |
|  | 5 | 2.15 | 31.2 |  | 2.15 | 31.2 |
|  | 6 | 5.51 (dd, 14.8, 9.0) | 131.7 |  | 5.51 (dd, 14.8, 9.0) | 131.7 |
|  | 7 | 6.02 (d, 14.8) | 130.7 |  | 6.02 (d, 14.8) | 130.7 |
|  | 8 | 6.04 (d, 14.8) | 130.7 |  | 6.04 (d, 14.8) | 130.7 |
|  | 9 | 5.70 (dd, 14.8, 9.0) | 134.9 |  | 5.70 (dd, 14.8, 9.0) | 134.9 |
|  | 10 | 2.14 | 32.0 |  | 2.14 (m) | 32.0 |
|  | 11 | 2.16 | 31.2 |  | 2.16 (m) | 31.2 |
|  | 12 | 5.55 (dd, 14.8, 9.0) | 131.7 |  | 5.55 (dd, 14.8, 9.0) | 131.7 |
|  | 13 | 6.03 (d, 14.8) | 130.7 |  | 6.03 (d, 14.8) | 130.7 |
|  | 14 | 6.14 (d, 10.9) | 130.2 |  | 6.14 (d, 10.9) | 130.2 |
|  | 15 | 6.14 (d, 10.9) | 132.7 |  | 6.14 (d, 10.9) | 132.7 |
|  | 16 | 6.16 (d, 14.8) | 134.6 |  | 6.16 (d, 14.8) | 134.6 |
|  | 17 | 5.57 (dd, 14.8, 9.0) | 126.8 | 3.30 (C19-acyl chain) | 5.57 (dd, 14.8, 9.0) | 126.8 |
|  | 18a | 2.34 | 37.4 |  | 2.60 (m) | 37.4 |
|  | 18b |  |  |  | 2.34 (m) |  |
|  | 19 | 3.30 | 47.9 |  | 3.54 | 52.7 |
|  | 20 | 1.22 (d, 6.8) | 17.3 |  | 1.27 (d, 6.8) | 17.3 |
|  | 2-Me | 1.76 (s) | 12.1 | 2.21 (C4-acyl chain);<br>7.01 (NH-Ile_1) | 1.76 (s) | 12.0 |

Heteronuclear single quantum coherence spectroscopy (HSQC) experiments revealed several cross peaks between 5.5-6.5 ppm (H) and 126-136 ppm (C). With the help of homonuclear COrrelation SpectroscopY (COSY) and TOtal Correlated SpectroscopY (TOCSY) experiments, these signals were associated with a C_20_ polyketide chain containing six double bonds: starting from the acyl group (C1), we could identify a methylated double bond (C2-C3 methylated in C2), two methylenes (C4-C5), two conjugated double bonds (C6 to C9), two further methylenes (C10 -C11), a triene system (C12 to C17) and a methylene (C18) (Fig. S8). The proton at 3.30 ppm, belonging to the C19 methine, showed COSY correlation with the terminal methyl group (C20) at 1.22 ppm and with the C18 methylene group at 2.34 ppm. The chemical shift of the corresponding carbon (47.9 ppm, C19) was compatible with an amine functionalization. We also observed eight ^3^J_H,H_ coupling constants of about 15 Hz and two of 11 Hz (Table 2). This indicated an *E* stereochemistry at all double bonds except for the one at C14-C15, which has a *Z* configuration. This was confirmed by a J-RESolved Spectroscopy (JRES) NMR experiment (Fig. S9). The methyl at C2 shows a strong NOE cross peak with the Ile amide at 7.01 and with the methylene in C4 (Fig. S8), which suggests an *E* configuration of the C2-C3 double bond, with the methyl at C2 and the proton at C3 on the opposite side of the double bond. The configuration of C19 could not be assigned from spectroscopic data. However, biosynthetic considerations (see below) suggest an *S* configuration. Overall, these analyses established the structure of the polyketide portion of allopeptimicins as a C_20_ chain carrying an amino group at C19 and three unsaturation systems (Fig.2).

Elucidanting the structure of the peptide portion required substantially more effort. In the 0-5.5 ppm portion of the proton spectrum, we could recognize spin systems belonging to one isoleucine (or *allo*-Ile; see below), one valine, two threonines, an *N*-methyl-leucine and two piperazic acids (Piz). A spin system containing a C2 carbon at 73.3 ppm (*δ*_C_) was consistent with an α-hydroxy acid, and COSY and TOCSY correlations demonstrated that it was part of a 2-hydroxy,4-methyl-glutaric acid (Hmg) moiety. The C3 carbon of one threonine was at 70.9 ppm instead of the standard 67 ppm, suggesting that this carbon was constrained and probably involved in peptide cyclization. An olefinic CH signal at 6.96/145.6 ppm, which was not associated with the polyketide chain, was included in one of the Piz spin systems, indicating an unsaturation between C5 and N5.

After adding H_2_O to the NMR sample, amide signals appeared and, with a combination of Rotating Frame Overhauser Enhancement Spectroscopy (ROESY) and Nuclear Overhauser Effect Spectroscopy (NOESY) experiments, we were able to connect few amino acids (Fig. S8). In particular, the amide signal at 7.00 ppm in the Ile spin system shows a correlation with both the methyl signal at 1.76 ppm (at C2 of the acyl chain) and the olefinic proton at 6.25 ppm (at C3 of the acyl chain), suggesting that Ile (or allo-Ile) is amidated with the C_20_ acyl chain. Moreover, we observed NOE correlations between the Val amide at 7.58 ppm and the proton at 4.34 ppm (CH at C2 of Ile or *allo*-Ile) and between the amide at 7.76 ppm in the threonine with a constrained C3 and the signal at 4.16 ppm (CH at C2 of Val). An additional correlation was seen between the amide signal at 7.73 ppm of the other threonine and the signal at 4.99 ppm (CH at C2 of Hmg). These observations established the partial sequences acyl chain-Ile(allo-Ile)-Val-Thr and Hmg-Thr in the peptide.

As mentioned above, when allopeptimicins were exposed to alkaline conditions, all congeners released a fragment having *m/z* 391 [M+H]^+^, together with a variable larger portion (Fig. S2A and B). When a 4-mg sample of the A+B complex was exposed to those conditions, the larger hydrolytic fragments showed the tricuspid UV absorption and thus carried the triene moiety, while the small hydrolytic product showed little UV absorption. The latter was purified and analyzed by HR-MSMS and NMR (Fig. S10 and S11), providing a calculated molecular formula for *m/z* 391.2046 [M+H]^+^ of C_17_H_31_N_2_O_8_ (Table 1). 1D- and 2D-NMR experiments demonstrated the presence of five methyl groups (one on a nitrogen), two methylene groups and six methines (two on oxygens). It was possible to recognize the spin systems of Hmg, Thr and *N*-MeLeu, in accordance with the molecular formula and with the fragments observed by HR-MSMS (Fig. S10). In particular, the fragment at *m/z* 146.1158 [M+H]^+^ corresponding to a C_7_H_16_NO^+^ is consistent with a hydrated *N*-methyl leucine, suggesting that this amino acid is at the C-terminal position of the 391-amu fragment. Overall, these data are consistent with the hydrolytic fragment having the sequence Hmg-Thr-MeLeu and being released by hydrolysis of two ester bonds. The only way to fit the smaller hydrolytic fragment is by connecting the Hmg α-hydroxyl and C-terminus to a carboxyl group and a threonine hydroxyl, respectively, in the larger hydrolytic fragments, establishing the peptide sequence as Ile*(allo*-Ile)-Val-Thr-Piz-Piz-Hmg-Thr-MeLeu.

By deeply analyzing the NMR data acquired for the A congeners mixture, we observed that the protons related to the unsaturated Piz spin system integrated to lower values than those of the other Piz residue, suggesting that the 2-amu difference and one lesser acetylation site of congeners A2 and A4 with respect to A1 and A3 are due to the unsaturation at C5-N5 of one of the Piz moieties. Similarly, the isoleucine-related signals appeared to be weaker than those of the other amino acids, while the valine spin system seemed to be spitted in two. This phenomenon could be reasonably associated with the A3 and A4 congeners having an isoleucine as the first amino acid followed by a valine, while two consecutive valines are present in A1 and A2.

With insights from the hydrolytic fragment and the NMR analysis of the A congeners, we were able to interpret the HRMS/MS data (Fig. S12 and S13). In particular, the most intense fragments *m/z* 613.3173 [M+H]^+^ and *m/z* 847.5410 [M+H]^+^ are consistent with the two consecutive Piz located between acyl chain-Ile(*allo*-Ile)-Val-Thr and Hmg-Thr-MeLeu moieties. Moreover, comparison between the fragments originating from A1 and A3 shows that the extra methylene difference must be in the N terminal amino acid, compatible with the first amino acid being a Val in A1 and an isoleucine in A3. In addition, the fragmentations of A1 and A2 indicates that the unsaturation is present on Piz5.

For establishing the stereochemistry of the amino-acid building blocks, we resorted to Marfey’s method.^17^ Both threonine residues were found to be D-*allo*-Thr while valine showed was the L-configuration. Unfortunately, we were not able to distinguish L-Ile from L-*allo*-Ile (Fig. S14) and, due to the lack of adequate standards, we could not assign the stereochemistry to the remaining stereocenters: one in each of the piperazic acid moieties and two in Hmg moiety. However, analysis of the allopeptimicin BGC suggested the possible stereochemistry of the remaining α-carbons and pointed to *allo*-Ile as the first amino acid residue.

Extensive 1D- and 2D-NMR experiments were also carried out on a sample containing a mixture of allopeptimicin B congeners, with minor amounts of the A congeners (Table 2). The resulting spectra were almost identical to those obtained from the A congeners, except for a shift in few HSQC cross signals: the C19 proton and carbon signals shifted from 3.30 and 47.9 ppm to 3.54 and 52.7 ppm, respectively; the two C18 methylene protons (both at 2.34 ppm) were split in two different signals at 2.34 and 2.60 ppm, while the C18 carbon at 37.4 ppm remained unchanged (Fig. S15A); and the protons of the C20 methyl group (at 1.22 in the A congeners) were shifted to 1.27 (see Fig. S15B which reports the overlap of TOCSY experiments from the two datasets and highlights the C18-C20 spin system in B). These changes, together with the mass spectrometry evidence of the presence of a -SO_3_H moiety and the absence of one acetylatable group in the B congeners, are compatible with the installation of a sulfamic acid on the primary amine at C19 (Fig. 2). Unfortunately, the fragmentation of the B congeners was not very informative, since the sulfamate moiety was split giving the mass of the corresponding A congener (data not shown).

**Fig. 2.**
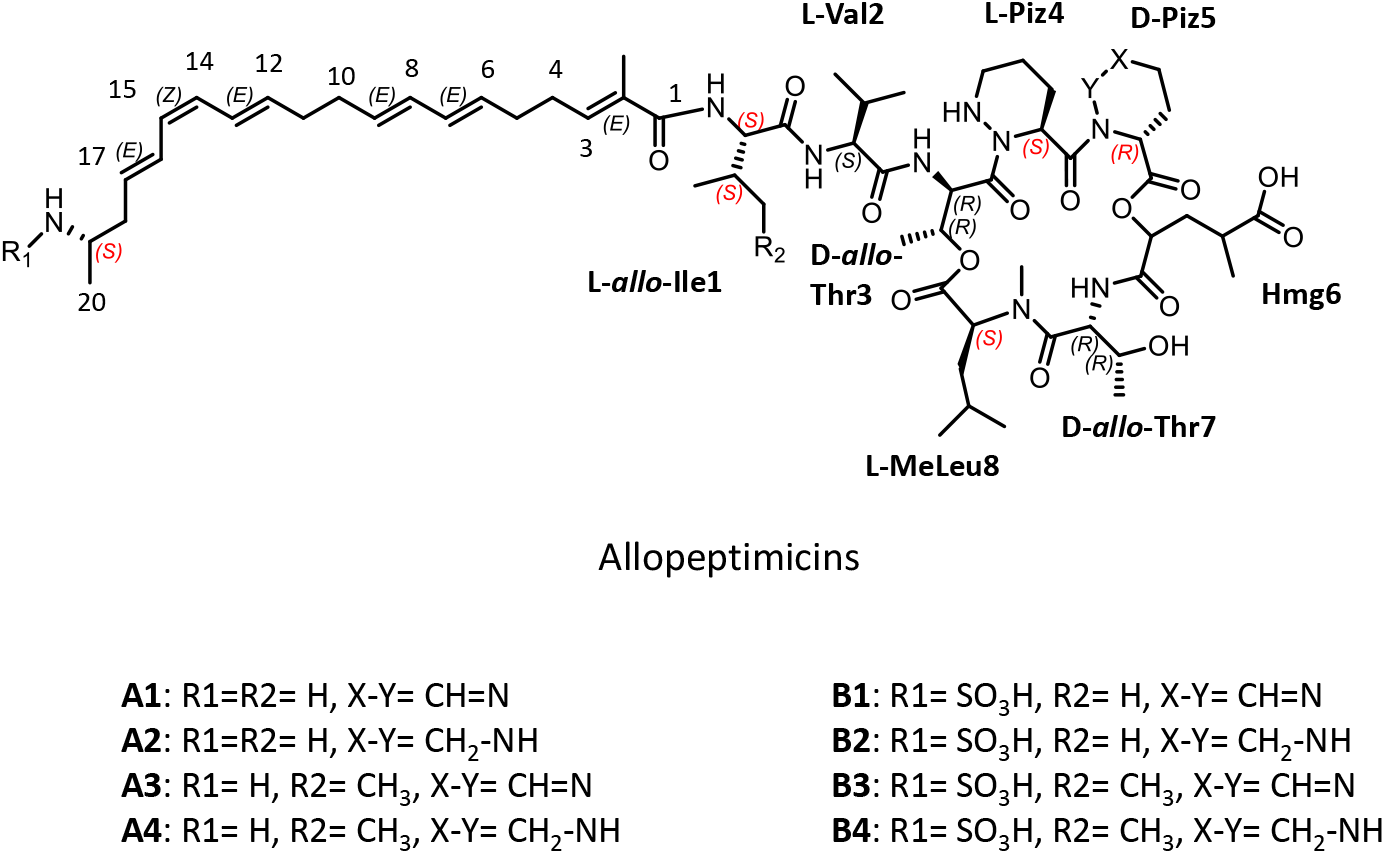
Structure of allopeptimicin congeners. In red stereocenters configuration inferred by bioinformatic analysis are reported.

In summary, the structures of allopeptimicins were established to be as reported in Fig. 2, with the B molecules arise from the cognate A congeners after installation of a sulfamate.

### The allopeptimicin biosynthetic gene cluster (BGC)

We readily identified the candidate *apt* BGC from a draft genome sequence of *Actinoallomurus* sp. ID145808 by using antiSMASH platform (https://antismash.secondarymetabolites.org/). The *apt* BGC is predicted to span 112 Kbp and consists of 35 CDSs associated with allopeptimicin biosynthesis, regulation and resistance. To most of those CDSs we were able to assign a role on the basis of homology searches and to establish the probable boundaries of the *apt* cluster (Table 3; Fig. 3A).

**Table 3.**
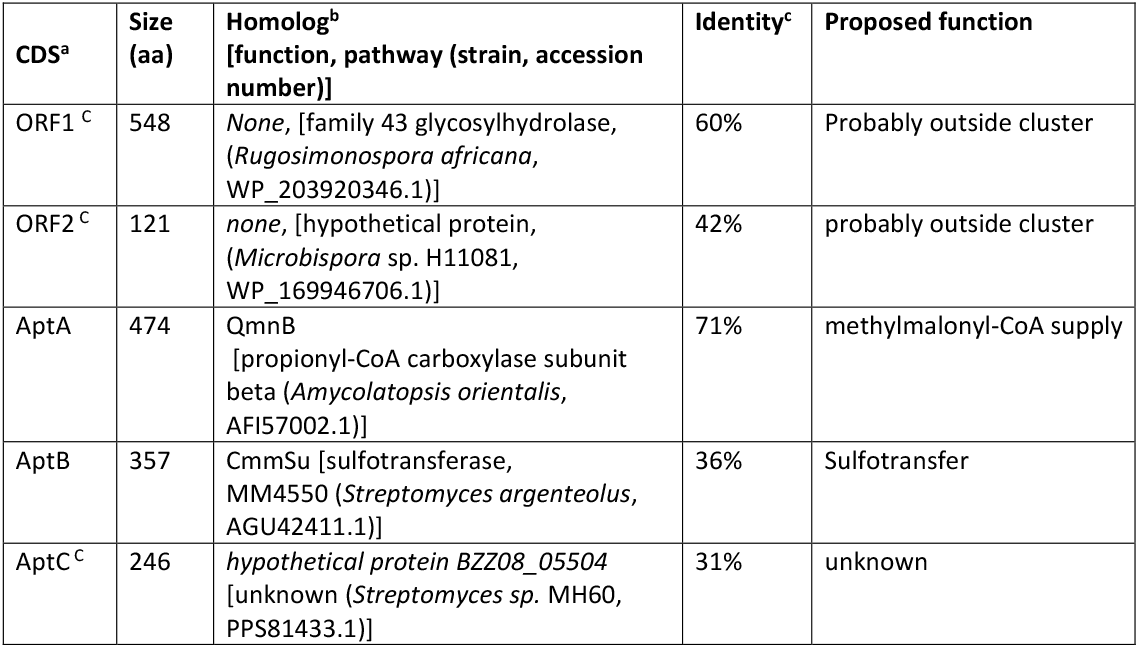
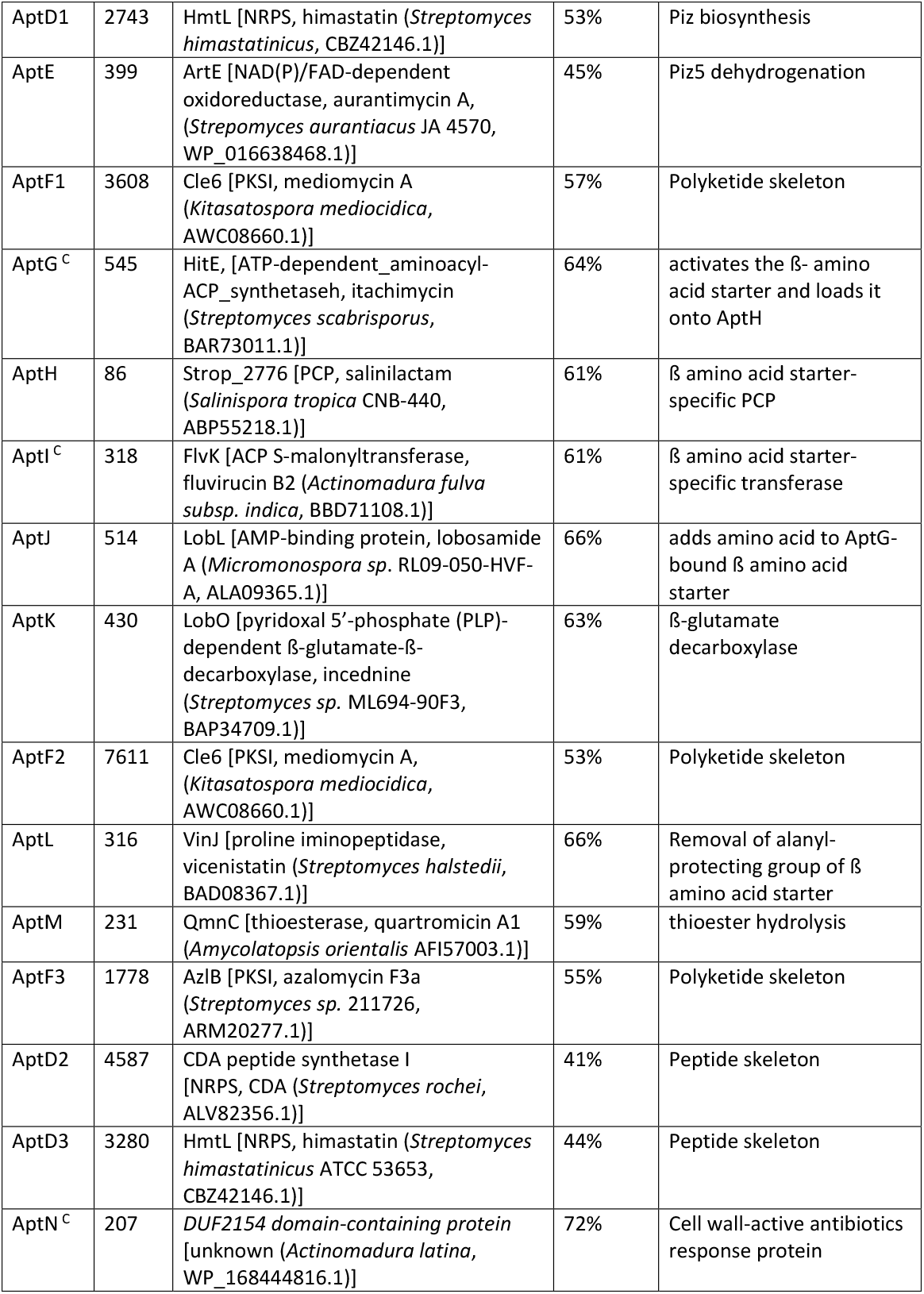
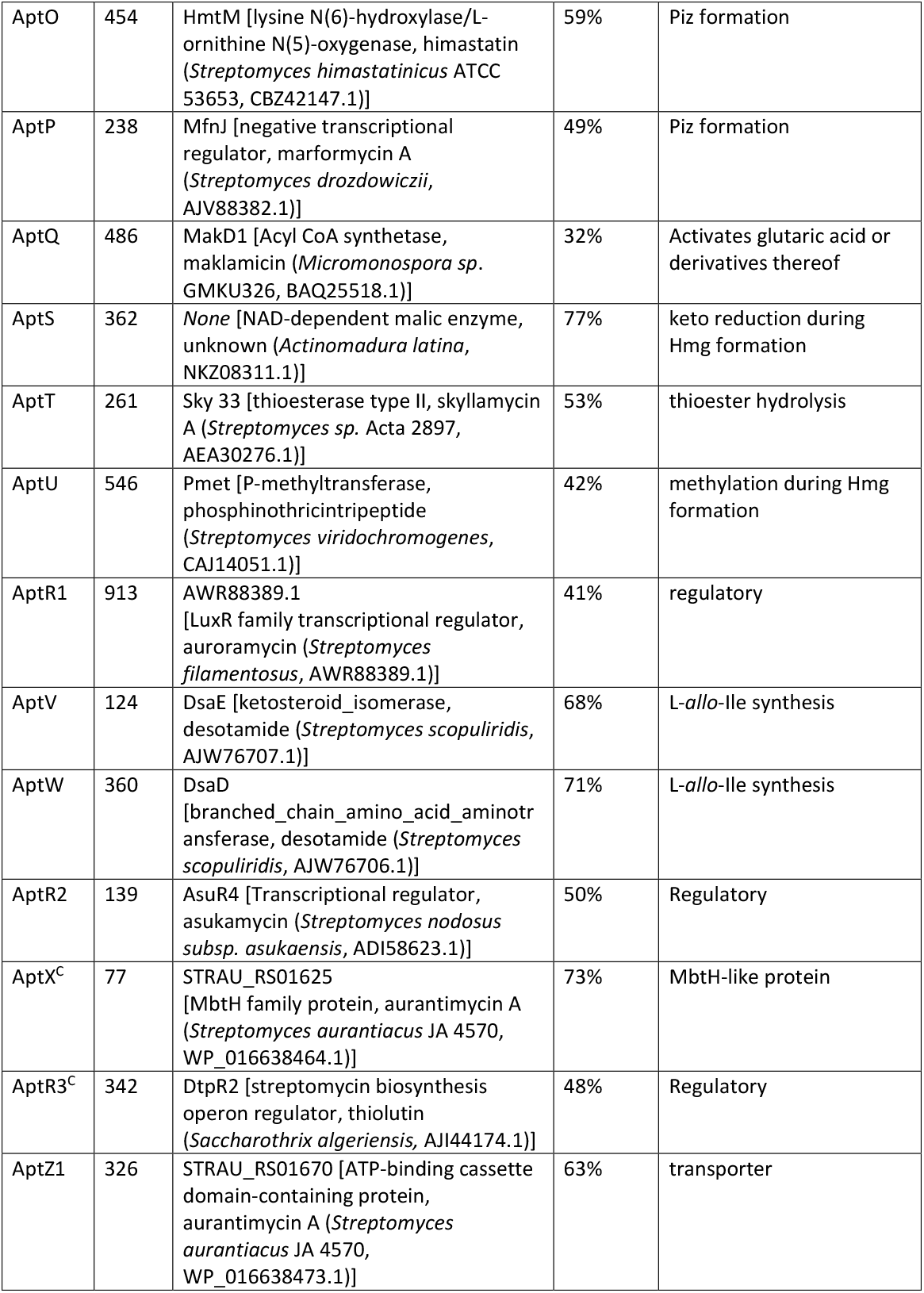
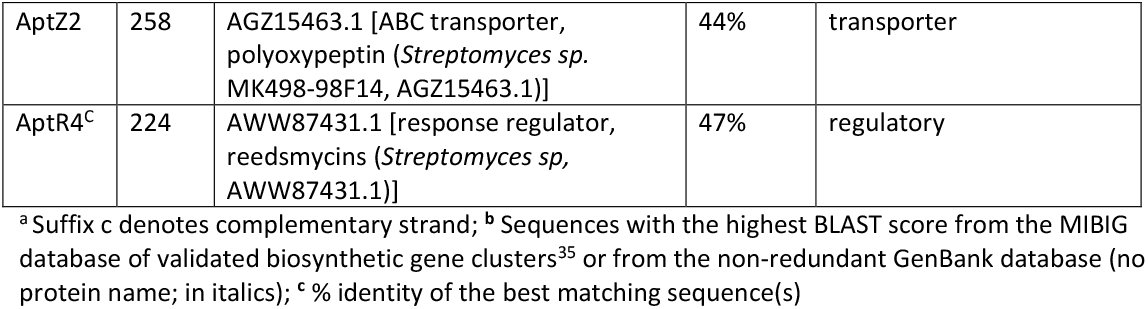
*apt* biosynthetic gene cluster and deduced function of each CDSs.

**Fig. 3.**
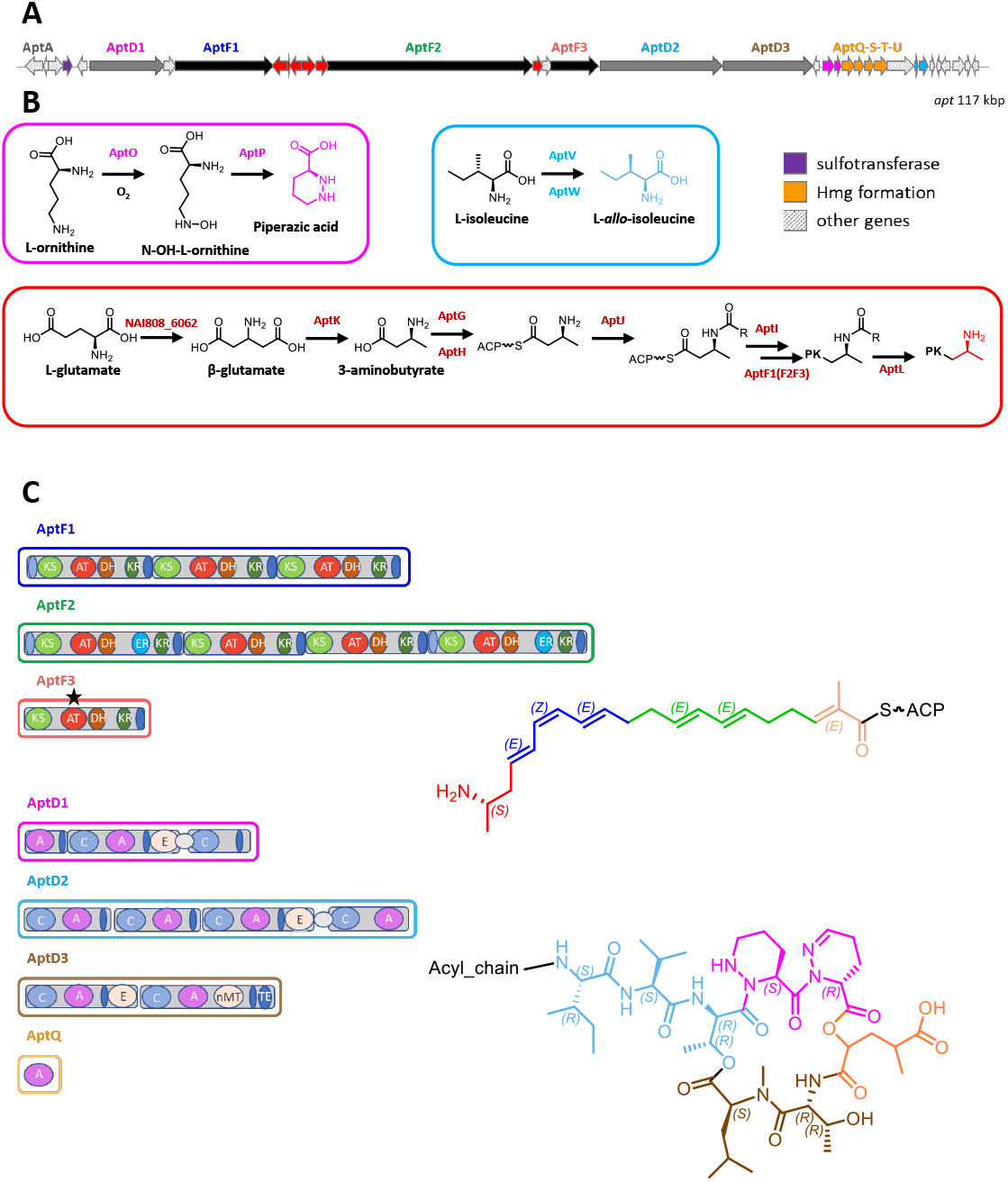
A) The allopeptimicin biosynthetic gene cluster, *apt*. The predicted functions for the open reading frames within the cluster are color-coordinated with the different panels. B) Proposed biosynthetic pathway for piperazic acid (Piz) (pink rectangle), L-*allo*-Ile (light blue rectangle) and the β-amino acid starter unit (red rectangle) with the individual steps catalyzed by *apt* enzymes. C) Domain arrangement and predicted biosynthetic intermediate for each PKS and NRPS polypeptide in the cluster, color-coordinated with the chemical structure. The black star indicates the AT domain recognizing a methylmalonate unit.

Consistent with the chemical structure, the *apt* BGC was found to encode three large polyketide synthases (PKS) for a total of eight modules, three non-ribosomal peptide synthetases (NRPSs) for additional eight modules, the gene products needed for the synthesis of four specialized precursors and for conversion of the A into B congeners (Table 3; Fig. 3A). In addition, one end of the BGC encodes regulators and transporters.

### Biosynthesis of the polyketide chain

The linear acyl chain of allopeptimicins contains an amine at C19, which would be consistent with a nitrogen-containing starter unit for the polyketide chain. Remarkably, the *apt* BGC contains a five-gene cassette (*aptGHIJK*) and an additional gene (*aptL*) with strong matches to genes coding for enzymes (Table 3; Fig. 3AB) involved in the formation and processing of the β-amino acid starter unit during macrolactam biosynthesis.^18^ After formation of the β-amino acid starter unit (see below), a stand-alone adenylation (A) domain selectively recognizes it and loads it onto a stand-alone peptidyl carrier protein (PCP). The amino group is then protected with L-alanine by another ATP-dependent ligase, followed by transfer onto the PKS starting module by a dedicated transferase. The alanyl-protecting group is eventually removed by a peptidase during or after polyketide elongation.^18,19^ AptG, AptH, AptI, AptJ and AptL are highly related (61 to 66% identity) to stand-alone A domains, PCPs, transferases, ATP-dependent ligases and oligopeptidases, respectively, from macrolactam BGCs (Table 3). In addition, AptK shows 63% identity to the PLP-dependent β-glutamate-β-decarboxylase from the incednine BGC.^20^ Of note, it has been demonstrated in macrolactams that the specificity for selecting the appropriate starter unit is due to the size of the amino acid side chain present at a critical position in the stand-alone A domain (position 220 in IdnL1).^21^ The presence of Leu220 in AptG as in IdnL1 suggests (*S*)-3-aminobutyrate is the starter unit for allopeptimicin biosynthesis and that this configuration is retained at C19. By analogy with the incednine pathway, we speculate that β-glutamate originates from L-glutamate through the action of a glutamate-2,3-aminomutase.^21^ While the apt BGC does not encode such an enzyme, a protein with 74% identity to the IdnL4 glutamate-2,3-aminomutase is encoded elsewhere by the ID145808 chromosome (unpublished data). Overall, these data allow us to propose a pathway for polyketide chain initiation from a β-amino acid starter (Fig. 3B). Preceding the *aptGHIJK* cassette is *aptF1*, encoding a PKS starting with an ACP domain and followed by three modules. Each module contains a dehydratase (DH) domain and all AT domains are predicted to be specific for malonate, based on computationally sequence similarity. These features make AptF1 a likely candidate for elongating the 3-amino butyrate starter (transferred onto the *N*-terminal PCP by AptI) through three malonate additions, followed by double bond formation after reduction and dehydration.

The C_10_ intermediate polyketide chain formed by AptF1 would then be elongated by AptF2, which consists of four modules: based on sequence analysis all contain AT domains predicted to be specific for malonate, KR domains and DH domains, but only the first and last module possess ER domains, consistent with the presence of saturated bonds at C4-C5 and at C10-C11 in the final product and with two conjugated double bonds (Fig. 2 and Fig. 3C). The mono-modular AptF3 would then complete polyketide synthesis through the incorporation of a (2*S*)-methylmalonyl-CoA unit, as for the predicted specificity of its AT domain, and installation of a double bond by the consecutive action of a B-type KR domain^22^ and by a DH domain. No bioinformatic prediction could be made about the stereo-selectively of the other KR domains, as no consensus residues for A and B type KR could be identified^22^.

Overall, the number of PKS modules, their domain arrangements and the predicted (stereo)specificities are consistent with collinearity between PKS modules and the structure of the C_20_ polyketide chain. Both AptF1 and AptF2 are mostly related to Cle6 (modules 1-3), a PKS involved in the synthesis of the giant linear polyketide mediomycin (Table 3), which carries an amino group at the ω position;^23^ while AptF3 is mostly related to module 1 from AzlB, a PKS involved in the formation of azalomycin.^24^

### Peptide and amino acid synthesis

The eight *apt* NRPS modules are also organized in three polypeptides: AptD1, AptD2 and AptD3 (Fig. 3A). The likely candidate for elongating the polyketide chain is the tetra-modular AptD2, which consists of two C-A-T modules, followed by a C-A-T-E module and terminating with C-A. The predicted specificities of the first three A domains are Val, Val and Thr, which nicely matches the *N*-terminal portion of the octapeptide: Val(L-*allo*-Ile)-Val-Thr (Fig. 3C). In addition, the E domain present in the third module fits with the experimental observation of both threonines being in the D-*allo* configuration (Fig. S14). The amino acid incorporated by the fourth module could not be predicted bioinformatically. The octapeptide chain is likely to be terminated by AptD3, a bimodular NRPS consisting of C-A-T-E and C-A-MT-T-TE domains, with predicted specificities for Thr and Leu. This nicely fits with Thr7 epimerization at the α-carbon, *N*-methylation of Leu8 and TE-catalyzed ester formation between Leu8 and Thr3 to close the macrocycle (Fig. 3C).

By exclusion, the remaining three residues (Piz4, Piz5 and Hmg6) are likely to be incorporated by the fourth module of AptD2 and by the bi-modular AptD1 that, overall, account for the expected three C, A and T domains, plus one E domain, but with an unusual arrangement (Fig. 3C): the second module in AptD1 lacks an A domain, which is present at the C-terminus of AptD2. The unusual domain organization of the second AptD1 module (C-T) has been observed also in HmtI and in KtzE from the himastatin and kutznerides BGCs, respectively: in both cases it has been suggested that the C domain forms an ester linkage between the amino acid loaded onto the upstream T domain and a hydroxy-acid acceptor.^25,26^ The same may happen during allopeptimicin biosynthesis, with the second C domain in AptD1 forming an ester linkage between Piz and Hmg. The latter could either be activated by the last A domain of AptD1 or provided in trans by the free-standing A domain AptQ (see below). While the amino acids incorporated by the two A domain in AptD1 could not be bioinformatically predicted, they share 95% identity, suggesting they recognize the same amino acid, hence Piz. In summary, we could tentatively propose the peptide segments recognized by the different modules as shown in Fig. 3C, although experimental work will be necessary to establish the role of the individual domains.

The E domains present in the third and first module of AptD2 and AptD3, respectively, directing the epimerization are consistent with the presence of two *D*-configured threonine residues at positions 3 and 7. If the central module of AptD1 is involved in adding the second Piz residue to the peptide chain, the associated E domain would then epimerize Piz5. Assuming that Hmg is in the L-configuration, the cyclic peptide would consist of alternating L- and D-amino acids, a feature often found in many NRPS-made peptides.^27^

Synthesis of the non-proteinogenic amino acids L-*allo*-Ile and Piz can be accounted for by the gene cassettes *aptOP* and *aptVW*, respectively (Fig. 3A and B). In the desotamide pathway, L-*allo*-Ile arises from *L*-Ile through the action of the pyridoxal 5’-phosphate-linked aminotransferase DsaD and the isomerase DsaE.^28^ Consistently, AptV and AptW show 64 and 70% identity with DsaD and DsaE, respectively (Table 3). In the kutznerides pathway, Piz is formed from L-ornithine through the consecutive action of the FAD-dependent *N*-hydroxylase KtzI and the heme-dependent dehydratase KtzT.^26,29^ AptO and AptP display 58 and 49% identity with KtzI and KtzT, respectively (Table 3). The unsaturation in Piz5 could result from the action of AptE, which shows 45% identity with ArtE, a FAD-dependent oxidoreductase from the aurantimycin BGC. Aurantimycin contains two Piz residues and the complex includes congeners with unsaturated Piz moieties.^30^ It remains to be established whether AptE acts on free Piz or after peptide synthesis.

While to our knowledge there is no description of a pathway to Hmg formation, the *aptQSTU* cassette, consisting of closely linked and likely translationally coupled genes, represents a likely candidate for it. AptS is a NADP-dependent oxidoreductase with 38-77% identity to NAD-dependent malic enzymes, and thus may be involved in converting 2-ketoglutarate in the corresponding hydroxy form. AptU is a radical SAM methyltransferase, likely to install the 4-methyl group. The free-standing A domain AptQ might be involved in activating 2-ketoglutarate before further modifications or it might interact in trans with AptD1 (see above). A role for the free-standing thioesterase AptT remains to be established. Further work will be necessary to establish the individual roles of enzymes for Hmg biosynthesis.

### Resistance determinants

The allopeptimicin B congeners carry a sulfate linked to the amine at C19 of the alkyl chain (Fig. 2). Close to the left-hand end of the *apt* BGC lies *aptB*, encoding a protein with 36% identity with CmmSu (Table 3), the enzyme catalyzing *O*-sulfonation of the C6 side chain during biosynthesis of the carbapenem MM4550 in *Streptomyces argenteolus*.^31^ Given the observed sequence identity, AptB is likely involved in formation of an N-S bond, thus transforming each A congener into the corresponding B form. Since the B congeners are devoid of activity (see also below), AptB is likely a self-protecting mechanism in *Actinoallomurus* sp. ID145808. An additional resistance determinant might be AptN, which shows 91% identity to members of DUF2154, a domain found in proteins responding to cell wall-active antibiotics in staphylococci, including YvqF, which is associated with glycopeptide resistance.^32^

### Additional genes

Additional functions related to allopeptimicin biosynthesis are: AptX, a member of the MbtH family, small proteins often associated with NRPSs;^33^ AptA, highly related to the β subunit of propionyl-CoA carboxylase, and thus a potential source of methylmalonyl-CoA for the last PKS extension step; and the additional free-standing TE AptM, a feature often observed in PKS and NRPS BGCs.^34^ Finally, the *apt* BGC contains *aptC* as a single gene of unknown function (Table 3).

While the left-hand end of the cluster is likely represented by *aptA*, the other end contains several regulatory and export genes: *aptR1*, encoding a member of the LuxR family of transcriptional regulators, and *aptR2*, with high similarity to a regulator from the asukamycin BGC, are located upstream and downstream, respectively, of the *aptVW* L-*allo*-Ile synthesis cassette. After *aptX* there are genes encoding AptR3, another regulator of the LuxR family, and the ABC transporters AptZ1 and AptZ2 (Table 3).

### Bioactivity

Allopeptimicins A were found to be particularly active against staphylococci, streptococci and *Clostridioides difficile*, with MICs of ≤0.06-0.125 µg/mL. Activity was also detected against other Gram-positive bacteria, such as *Bacillus subtilis, Micrococcus luteus* and enterococci (Table 4). Except for some activity against *Helicobacter pylori* (MIC 16 µg/mL), no activity was observed against other Gram-negative species (Table 4). No activity was observed against eukaryotic microorganisms (data not shown). In contrast, allopeptimicins B were totally inactive (Table 4). In time-kill experiments with exponentially growing *S. aureus* ATCC 6538P, allopeptimicins A at 32 µg/mL effectively killed the bacteria within 3 hours (Fig. S16A).

**Table 4.** Antimicrobial activities of allopeptimicins A and B against selected bacterial species.

| strain<sup>a</sup> | MIC ($\mu\text{g/mL}$) | | |
| --- | --- | --- | --- |
|  | Allopeptimicin A | Allopeptimicin B | VAN<sup>b</sup> |
| *Staphylococcus aureus*<br>ATCC6538P | 0.125 | >128 | 0.25 |
| *S. aureus* L3864 (MRSA) | 0.125 | >128 | 0.5 |
| *S. epidermidis*<br>ATCC12228 | 0.125 | >128 | 1 |
| *Streptococcus pneumoniae* L44 | ≤0.06 | >128 | 0.25 |
| *S. pyogenes* L49 | ≤0.06 | >128 | 0.25 |
| *Micrococcus luteus*<br>ATCC10240 | 1 | >128 | 0.125 |
| *Bacillus subtilis* 168<br>ATCC27370 | 0.125 | >128 | 0.25 |
| *Enterococcus faecium*<br>L568 | 8 | >128 | 1 |
| *E. faecalis* L559 | 8 | >128 | 4 |
| *Clostridioides difficile* 421<br>L4013 | ≤0.06 | nt | 1 |
| *Helicobacter pylori* L4219 | 16 | nt | >128 |
| *Pseudomonas aeruginosa*<br>ATCC27853 | >128 | >128 | >128 |
| *Escherichia coli*<br>ATCC25922 | >128 | >128 | >128 |

<sup>a</sup>Strains designated with an L prefix are clinical isolates (collected in Italy or USA) from the NAICONS collection. Other strains are from public collections as indicated. American Type Culture Collection, USA. <sup>b</sup> Vancomycin (Van) used as control antibiotic.

We then investigated the effect of allopeptimicins A on macromolecular synthesis in *S. aureus* cells by monitoring the incorporation of labeled precursors (5-[^3^H] thymidine for DNA, [^3^H] uridine for RNA, L-[^3^H] tryptophan for protein and [^3^H] glucosamine for cell wall synthesis: we readily observed that cell wall biosynthesis was inhibited by 50% already at 1.6 µg/ml allopeptimicins A, while the concentrations inhibiting by 50% (IC50) were 5.7 µg/ml for DNA synthesis, 35 µg/ml for RNA synthesis and >50 µg/ml for protein synthesis, thus suggesting that cell wall synthesis is the primary target for this antibiotic (Fig. S16B). Consistently, when tested against different *S. aureus* mutants selected through serial passages, phenotypically and genotypically related to vancomycin insensitive *S. aureus* (VISA) strains,^36^ we observed an increase in the MIC of allopeptimicins A (Fig. S16c). Against these same mutants, a similar shift in MICs was observed for other *bona fide* antibiotics that bind to the essential peptidoglycan precursor Lipid II such as vancomycin, ramoplanin and NAI-107.^36^

### Self-defense mechanism in the producer strain *Actinoallomurus* sp. ID145808

The producer strain ID145808 was isolated from a soil sample collected in California (US). It was classified as *Actinoallomurus* on the basis of its 16S rRNA gene sequence, which shows 99% identity to the sequence from *Actinoallomurus bryophytorum*.

When ID145808 was grown in AF-AS medium, production of allopeptimicins started to be detectable at 72 h (Fig. 4A) and remained essentially constant up to 120 h. During this period, congeners A and B were produced in similar amounts. Starting at 144 h, the production rate increased, with the A congeners reaching 115 mg/L at 240 h, while production of allopeptimicins B increased at a lower rate, leading to a final ratio of about 75:25 A:B (Fig. 4A). Cultivation times longer than 240 h did not lead to increased production.cSupplementing the medium with 1 g/L L-threonine did not affect the early phases of production in terms of total allopeptimicins and A:B ratio (Fig. 4B). However, L-threonine supplementationcafforded higher titers at 240 h and extended the production phase up to 288-312 h, leading to 180 mg/L. L*-*threonine supplementation did not alter the A:B ratio, which remained 3:1 at long incubation times.

**Fig. 4.**
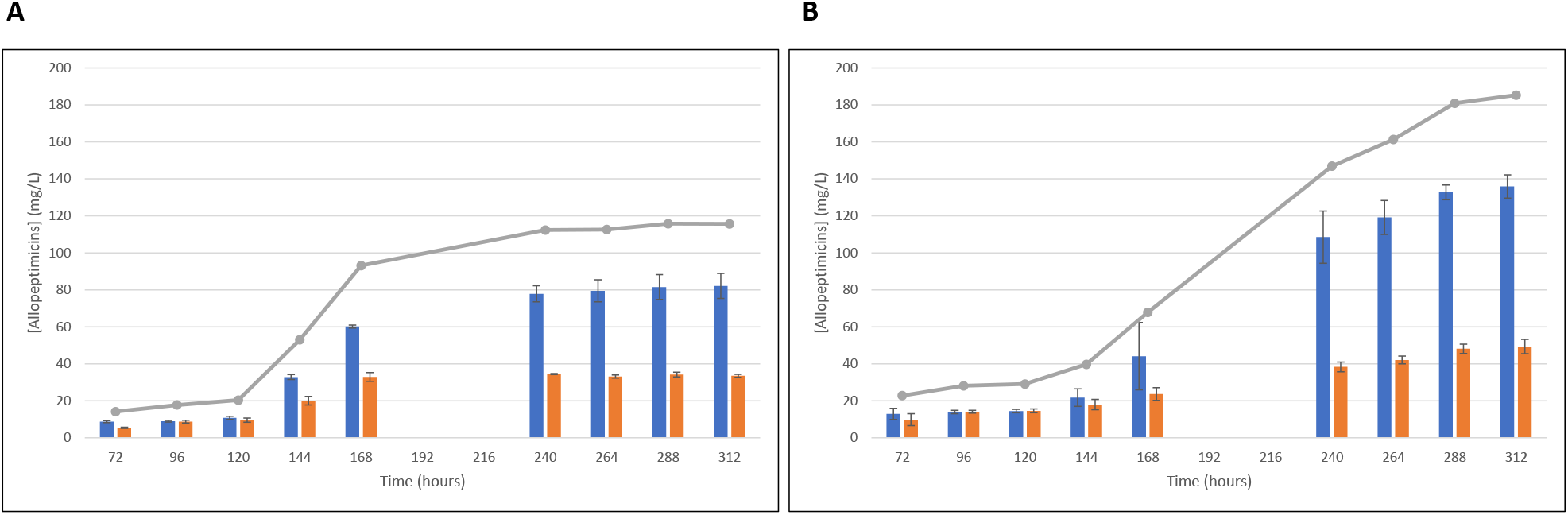
Production of allopeptimicins without (A) and with (B) threonine supplementation. Values calculated as means of three replicates with standard deviation. Allopeptimicins A and B are indicated by blue and orange bars, respectively. The grey line indicates the total complex (A+B) observed.

Since congeners A possess antibacterial activity while congeners B do not, the switch in congener ratio upon prolonged cultivation prompted us to analyze the sensitivity of *Actinoallomurus* ID145808 versus its own products (Fig. S17). The growth of the producer strain was not affected by spotting increasing amounts of a sample of allopeptimicins B (which contained around 5% of A congeners), until a slight growth disturbance was observed when 50 µg were spotted on the plate. Conversely, 2 µg of the A congeners fully prevented growth, indicating that *Actinoallomurus* ID145808 is sensitive to the A congeners when growing on agar plates, a condition when visible colonies appear after 72-96 h. We can thus hypothesize that formation of allopeptimicins B is a mechanism of self-protection. Since *Actinoallomurus* sp. ID145808 grows as large mycelial clumps in liquid medium, it is conceivable that non-dividing portions of a mycelial unit produce the active congeners while actively duplicating sectors express *aptB* to protect themselves. Further experiments need to be done to understand self-resistance in *Actinoallomurus* sp. ID145808.

## Conclusions

Allopeptimicins possess unusual structural features. Few examples exist of hybrid polyketide-peptides carrying such a long and unsaturated acyl chain: the closest relative may be the anabaenolysins from the cyanobacterium *Anabaena*, which consist of a C_18_ poly-unsaturated fatty acid that contributes three carbon atoms to a peptidic macrocycle.^37^ In addition, trienes with an *E-Z-E* configuration are rarely encountered: examples are represented by some colabomycin congeners^38^ from *Streptomyces*, and aurafuron B^39^ and spirangiens^40^ from myxobacteria. A sulfamate moiety has been observed only in metabolites from marine macroorganisms: some minalemine congeners from the tunicate *Didemnum rodriguesi*;^41^ scleritodermin from the lithistid sponge *Scleritoderma nodosum*;^42^ and some ianthesine congeners from the marine sponge *Ianthella* sp..^43^ To our knowledge, there is only one precedent for metabolite carrying an Hmg moiety: the D congeners of destruxins, hexadepsipeptides produced by *Metarhizium anisopliae*^44^ and other soil fungi. All the above features, along with the uncommon presence of two ester bonds in a single macrocycle, make allopeptimicins truly unique metabolites.

The structural uniqueness of allopeptimicins is reflected in a complex BGC, with dedicated gene cassettes devoted to formation of specialized precursors and modular assembly lines related to those from different BGCs. Thus, the *apt* BGC is one more example of a BGC with a mosaic structure, a feature common to many BGCs.^4,45^ Several aspects of the allopeptimicin NRPS remain obscure, including how the Piz-Piz-Hmg portion is assembled.

Structurally, allopeptimicins A consist of a relatively rigid macrocycle with a long acyl chain with few rotatable bonds and terminating with a positive charge. The available evidence - from macromolecular syntheses to reduced activity on VISA strains - suggest that allopeptimicins A interfere with cell wall biosynthesis. It is tempting to speculate that the macrocycle interacts with some essential cell wall components near the bacterial membrane, while the acyl chain embeds into the membrane, with further ionic interactions between the negatively charged phospholipids and the amino group at C-19. Turning the positive into a negative charge through sulfamidation would abolish this interaction with total loss of activity. Not many precedents exist of metabolites whose activity is modulated by sulfation: a notable example is represented by the paralytic shellfish saxitoxins, in which *O*- and/or *N*-sulfation leads to a reduced toxicity.^46^ In any case, turning a positive into a negative charge by formation of a sulfamate represents one more example of the ingenuity of soil bacteria in devising mechanisms for self-protection.

## Materials and Method

### General experimental procedures

^1^H and ^13^C 1D and 2D NMR spectra (COSY, TOCSY, NOESY, ROESY, HSQC, HMBC) were measured in CD_3_CN:D_2_O 8:2 with or without drops of H_2_O at 25 °C using a Bruker Avance II 300 MHz spectrometer. LC-MS/MS analyses were performed on a Dionex UltiMate 3000 HPLC system coupled with an LCQ Fleet (Thermo Scientific) mass spectrometer equipped with an electrospray interface (ESI) and a tridimensional ion trap. The column was an Atlantis T3 C-18 5 μm X 4.6 mm X 50 mm maintained at 40 °C at a flow rate of 0.8 mL/min. Phase A was 0.05% trifluoroacetic acid (TFA), phase B was 100% acetonitrile. The elution was executed with a 14 min multistep program that consisted of 10, 10, 95, 95, 10 and 10% phase B at 0, 1, 7, 12, 12.5 and 14 min, respectively. UV-Vis signals (190-600 nm) were acquired using the diode array detector. The *m/z* range was 110-2000 and the ESI conditions were as follows: spray voltage of 3500 V, capillary temperature of 275 °C, sheath gas flow rate at 35 units and auxiliary gas flow rate at 15 units. High resolution MS spectra were recorded at Unitech OMICs (University of Milano, Italy) using a Dionex Ultimate 3000 HPLC system coupled with a Triple TOF® 6600 (Sciex) equipped with an ESI source. The experiments were carried using the same column and eluting system as described for low resolution LC-MS analysis. The ESI parameters were the following: curtain gas 25 units, ion spray voltage floating 5500 v, temperature 50 °C, ion source gas1 10 units, ion source gas2 0 units, declustering potential 80 v, syringe flow rate 10 μL/min, accumulation time 1 sec.

### Bacterial strains, growth and fermentation procedures

For metabolite analyses, 88 *Actinoallomurus* strains from the NAICONS collection were propagated from cryovials on BTT and S1 agar plates^47^ and incubated at 30 °C for ca. 14 d. Mycelium from BTT plates was inoculated into 15 mL of AF-A medium^12^ in 50-mL Erlenmeyer flask and incubated 72 h at 30 °C in a rotary shaker at 200 rpm. Then, 10% seed culture was transferred into 15 mL AF-A medium in 50-mL Falcon tube supplied with 1 g of 5-mm glass beads. After 7 d in a rotary shaker at 250 rpm and at 30 °C, cultures were harvested. Two different extracts were prepared from each culture: one by solvent extraction of the pelleted mycelium and by solid-phase adsorption of the cleared broth essentially as described.^8^ The 176 extracts generated were analyzed individually by transferring 500 µL in HPLC vials prior to drying.

For allopeptimicin production, 2 ml of a frozen mycelium stock of strain ID145808 was inoculated into 25 ml of AF-A medium in a 300-mL Erlenmeyer flask and incubated 72 h at 30 °C in a rotary shaker at 200 rpm. Then, 6 ml was transferred into 100 mL AF-AS medium (AF-A supplemented with 10 g/L soluble starch) in 500-mL flasks supplied with 10 g of 5 mm glass beads. Where required, 1 g/L L-threonine (Sigma-Aldrich) was added from a filter-sterilized 50 mg/mL stock solutions. The flasks were incubated at 30 °C, on a rotary shaker at 200 rpm. For allopeptimicin analysis, 0.5 mL was withdrawn from the production culture and transferred into 2-mL Eppendorf tubes containing 0.5 mL MeCN. Samples were kept at 40 °C under constant shaking (800 rpm) for 1 h, then centrifuged for 3 min at 13 000 rpm. The supernatants were recovered and 500 µL were transferred into 1.5-mL glass vials for LC-MS/MS analysis.

### Processing of LC-MS/MS Data

.mzXML files from 176 *Actinoallomurus* extracts were processed with MZmine 2 v53^48^ as follows: (A) MassDetection = retention time, auto; MS1 noise level, 1E3; MS2 noise level, 0E0. (B) ADAP chromatogram builder = retention time, auto; MS-level, 1; min group size in no. of scans, 3; group intensity threshold, 5E1; min highest intensity, 1E1; *m/z* tolerance, 0.5 Da. (C) Chromatogram deconvolution = baseline-cutoff algorithm; min peak height, 1E2; peak duration, 0.03−1 min; baseline level, 1E2; *m/z* range MS2 pairing, 5 Da; RT range MS2 pairing, 0.25 min. (D) Isotopic peaks grouper = *m/z* tolerance, 1 Da; RT tolerance, 0.25 min; monotonic shape, no; maximum charge, 2; representative isotope, most intense. (E) Join peak alignment = *m/z* tolerance, 1 Da; RT tolerance, 0.25 min; weight for *m/z*, 30; weight for RT, 70. (F) MS Features with no accompanying MS2 data were excluded from the analysis. The resulting MS feature list was exported to the GNPS-compatible format, using the MS Feature-Based Molecular Networking (FBMN) workflow (version release_28.2) on GNPS.^5^

Parameters were adapted from the GNPS documentation as follows: MS2 spectra were filtered to remove all MS/MS fragment ions within ±17 Da of the precursor *m/z* value; only the top six fragment ions in the ±50 Da window through the spectrum were utilized, with a minimum fragment ion intensity of 50. The MS/MS fragment ion tolerance was set to 0.9 Da and the precursor ion mass tolerance was set to 2 Da. Edges of the created molecular network were filtered to have a cosine score above 0.5 and more than 2 matched peaks between the connected nodes, with a maximum of 10 neighbor nodes for one single node. The maximum size of clusters in the network was set to 50 and the maximum shift between precursors was set at 300 Da. The MS2 spectra in the molecular network were searched against GNPS spectral libraries. Reported matches between network and library spectra were required to have a score above 0.7 and at least 3 matched peaks. The molecular network was visualized using Cytoscape 3.9.0.^49^

### Allopeptimicin Purification

A 1-L, 312-h culture of strain ID145808 in AF-AS medium, containing 80 and 38 mg/L of A and B congeners, respectively, was centrifuged 10 min at 3500 rpm in 250-mL centrifuge bottles. The pellet was discarded while the cleared broth was pooled, brought to pH 4 with 5.5 mL 1M HCl and extracted with 2×500 mL dichloromethane:methanol 8:2 in a 2-L separating funnel. The organic phase was dried under reduced pressure and the powder was washed twice with 5 mL H_2_O by transferring the suspension into a 15-mL Falcon tube, centrifuging 10 min at 4000 rpm and discarding the washing solution. Then, the pellet was dissolved in 5 mL dichloromethane and resolved using a CombiFlash RF (Teledyne ISCO) medium pressure chromatography system on a 24 g RediSep RF Silica flash column (Teledyne ISCO). Flow was 20 mL/min, phase A was dichloromethane and phase B was methanol. The column was previously conditioned with 100% phase A for 7 min, followed by a 21-min gradient to 60% phase A and kept at 60% phase A for 10 min. Fractions of 15-mL were collected and analyzed by LC-MS. Fractions with similar purity were pooled and dried under vacuum, yielding about 20 mg of a mixture of A congeners and 7 mg of a mixture of B congeners, the latter containing about 15% w/w A congeners. These samples were used for NMR characterization and for the sensitivity of *Actinoallomurus* ID145808 versus its own products. For bioactivity studies the A congeners mixture was used as such while the B mixture was further purified as follows: 1.5 mg of the powder were dissolved in 400 µL MeCN 60% and chromatographed using 20 analytical HPLC runs on a Dionex UltiMate 3000 HPLC system, using the same conditions as described for LC-MS analysis and collecting the fractions at 7.8 minutes. About 1 mg of pure allopeptimicin B congeners were achieved.

### Chemical modifications of allopeptimicins

Reactions were performed on partially purified samples obtained from different fermentations containing a mixture of A and B congeners. **Hydrolysis**: A 4-mg sample (6:4 A:B) was dissolved in 1 mL 50% acetonitrile, then 1 mL 1M NaOH was added. After 1 h stirring at room temperature (RT), 0.5 mL H_2_O was added to the reaction mix. After LC-MS analysis, the reaction mix was buffered to pH 7 by adding 1M 0.5 mL HCl and resolved using a 5g flash C18 Isolute SPE cartridge (Biotage). After loading the sample, the column was washed with 10 mL H_2_O, then eluted with 10 mL 30% acetonitrile, yielding a fraction containing the smaller hydrolysis product (with *m/z* 391 [M + H]^+^), which was dried, dissolved in 0.5 mL D_2_O and analyzed by NMR. **Acetylation**: A 1-mg sample (6:4 A:B) was dissolved in 0.2 mL acetonitrile, then 0.2 mL acetic anhydride was added. The reaction was stirred 1 h at RT and then analyzed by LC-MS. **Amidation**: A 2-mg sample (4:6 A:B) was dissolved in 500 μL dry DMF and the pH adjusted to 8 by adding 2 μL methylamine. After adding 1 mg PyBOP, the mixture was stirred for 10 min at RT and analyzed by LC-MS. **Stereochemistry determination**: A sample of allopeptimicins (2 mg, 9:1 A:B) was treated as described,^17^ together with the following amino acid standards: L-Thr, D-Thr, L-*allo*-Thr, D-*allo*-Thr, L-Val, D-Val, L-Ile and L-*allo*-Ile, and then analyzed by LC-MS using a HiQ sil C18 HS column (4.6 × 250 mm, Kya Tech Corporation, Japan). Phase A was 0.05% trifluoroacetic acid, phase B was acetonitrile. The samples were run using a gradient from 10 to 90% B in 45 min at a flow rate of 0.8 mL/min.

### Antimicrobial assays

The antibacterial activity of extracts was determined as described.^16^ MICs were determined by micro-dilution according to the Clinical and Laboratory Standards Institute guidelines (https://clsi.org).^50^ All aerobic strains were grown in Müller Hinton broth containing 20 mg/L CaCl_2_ and 10 mg/L MgCl_2_ except for *Streptococcus* spp. and *Haemophilus influenzae*, which were grown in Todd Hewitt broth and Brain Hearth Infusion (with 0.5% yeast extract), respectively. Strains were inoculated at 1×10^5^ CFU/mL. For *Clostridioides difficile*, fresh cultures were diluted to 1×10^5^ CFU/mL with Brucella medium (BB) supplemented with hemin (5 μg/mL), vitamin K1 (1 μg/mL), lysed horse blood (5%) and oxyrase®(1:25 v/v), while for *Helicobacter pylori* the strain was streaked twice on Brucella agar plates, then colonies were resuspended to an OD_600_ of 0.8 and diluted 1:10 with BB containing 3% calf bovine serum. Bacteria were incubated at 37°C for 24 h except for *C. difficile* (48 h) and *H. pylori* (72 h). Allopeptimicins were dissolved at 10 mg/mL in dimethyl sulfoxide, while vancomycin, used as control, was dissolved in water. Samples were diluted with the test medium to the required concentrations.

Time-kill curves were performed as previously described^51^ using *S. aureus* ATCC 6538P. Growth kinetics was performed under the same conditions as MICs and recording OD_600_ every hour in a Synergy 2 reader (Biotek).

For evaluating the activity against strain ID145808, serial dilutions in 60% MeOH were prepared from stock solutions of allopeptimicins A (2 mg/mL) and B (5 mg/mL; this preparation contained 5% A congeners). Then, 10 µL of appropriate dilutions were spotted onto BTT agar plates, which had been previously seeded with strain ID145808. Positive and negative controls were 10 µg/mL apramycin and 60% MeOH, respectively. Plates were incubated at 30 °C for 10 days before scoring inhibition haloes.

### Macromolecular synthesis

The effect on macromolecular synthesis was tested by monitoring the incorporation of labeled precursors (5-[^3^H] thymidine for DNA, [^3^H] uridine for RNA, L-[^3^H] tryptophan for protein and [^3^H] glucosamine hydrochloride for cell wall synthesis as described.^52^

### Bioinformatic analyses

From a draft genome sequence of *Actinoallomurus* sp. ID145808, carried out using Illumina platform and MinION sequencing device (M.S., unpublished data), we identified the *apt* BGC using antiSMASH 6.0 at the default conditions.^53^ Then, the sequences of the *apt* genes were manually curated. BLAST analysis of individual CDSs was performed against the MIBiG 2.0 database of known gene clusters^35^ and against Protein Data Bank. The 16S rRNA gene sequence was analyzed following published procedures.^54^

The nucleotide sequences have been deposited in GenBank under accession numbers XXXX (apt BGC) and YYY (16S rRNA gene).

## Author contributions

M.I., S.D. and M.S. designed work, analyzed data and wrote the paper; A.G. performed strain cultivation, molecular networking analysis and analyzed data; M.I. and S.I.M. identified metabolites and analyzed data; M.I. purified metabolites and elucidated structures; C.B., P.L., A.G. and A.T. performed bioassays and analyzed data. All authors contributed to critical discussions, read and approved the final manuscript.

## Acknowledgments

This work was supported by grants to NAICONS from Italian Ministry of Research (No. DM60066) and to KtedoGen from the European Union’s Horizon 2020 research and innovation program under grant agreement No. 664588 (Nomorfilm). The authors thank Joao C. Cruz for early contributions to this project.

## Supplementary tables and figures

**Fig. S1**. Molecular network of 176 extracts from 88 *Actinoallomurus* strains, encompassing 725 features (=nodes). 384 features were organized in 52 molecular families. Pink nodes indicate features present in extract from *Actinoallomurus* ID145808 only. Enlargement shows selected interesting signals from ID145808.

**Fig. S2**. A) Extracted ion chromatograms of *m/z* 391, 851-868, 931-948, 1205-1222 and 1285-1302 [M+H]^+^ from LC-MS analysis of the starting allopeptimicin complex (black line) and the reaction with 0.5N NaOH for 1h (red line). B) MS/MS at 4.9, 6.3-6.5 and 7.0-7.4 min of the hydrolysis mixture. C) Extracted ion chromatograms of *m/z* 1205-1233 and 1285-1314 [M+H]^+^ from LC-MS analysis of the starting allopeptimicin complex (black line) and the reaction with 0.81 M CH3NH2 for 10 min (red line). D) MS/MS at 6.7-7.0 and 7.7-8.2 minutes from the amidation reaction. See Material section for the reaction conditions.

**Fig. S3**. ^1^H NMR (300 MHz, CD_3_CN:D_2_O:H_2_O 4:1:0.2) of allopeptimicins A.

**Fig. S4**. ^13^C NMR (75 MHz, CD_3_CN:D_2_O:H_2_O 4:1:0.2) of allopeptimicins A.

**Fig. S5**. ^1^H-^13^C HSQC NMR (300 MHz, CD_3_CN:D_2_O:H_2_O 4:1:0.2) of allopeptimicins A.

**Fig. S6**. Overlap of ^1^H-^13^C HSQC (blue/green) and ^1^H-^13^C HMBC (red) NMR (300 MHz, CD_3_CN:D_2_O:H_2_O 4:1:0.2) of allopeptimicins A.

**Fig. S7**. Overlap of ^1^H-COSY (red) and ^1^H-TOCSY (blue) NMR (300 MHz, CD_3_CN:D_2_O:H_2_O 4:1:0.2) of allopeptimicin A.

**Fig. S8**. Structure of allopeptimicin A3. COSY correlations are highlighted in blue. Red arrows show the main NOESY correlations.

**Fig. S9**. J-res NMR spectrum of allopeptimicins A. The red circle highlights the J-coupling relative to C14 and C15.

**Fig. S10**. HR-MSMS spectrum of the alkaline hydrolysis product with *m/z* 391.2046 [M+H]^+^ and proposed fragment assignment.

**Fig. S11**. NMR spectra of the alkaline hydrolysis product with *m/z* 391 [M+H]^+^(see text for details).

A) ^1^H NMR, B) TOCSY and C) overlap of ^1^H-^13^C HSQC (blue/green) and ^1^H-^13^C HMBC (red) acquired at 300 MHz in D_2_O. D) ^1^H and ^13^C NMR spectral data of the hydrolytic fragment of allopeptimicins.

**Fig. S12**. Characterization of allopeptimicin A1, A2 and A3 by HR-MSMS. A) HR-MS^2^ spectrum of 603.3622 [M+2H]^2+^, 604.3689 [M+2H]^2+^ and 610.3702 [M+2H]^2+^ B) HR-MS^2^ zoom of selected region.

**Fig. S13**. A) Table of found and calculated exact masses of fragments from allopeptimicin A1, A2 and A3; B) proposed assignments of observed MS fragments (PK corresponds to the polyketide chain).

**Fig. S14**. Extracted ion chromatograms of amino acid standards and allopeptimicins after treatment according to Marfey’s method.^17^ Extracted ion chromatograms of the threonines (A), valines (B) and isoleucines standards (C) aligned with the hydrolysis products of allopeptimicin.

**Fig. S15**. A) Overlap of a selected region of HSQC experiments of allopeptimicins A (blue/green) and allopeptimicins B (red/black). Arrows indicate the chemical shift of CH at position 19 and CH_2_ at position 18 in both samples. B) Overlap of a selected region of ^1^H-TOCSY experiments of allopeptimicins A (blue/green) and allopeptimicins B (red/black). The black rectangle highlights the C18 (2.60-2.30) - C19 (3.54) - C20 (1.27) spin system in allopeptimicin B. NMR experiments acquired at 300 MHz in CD_3_CN:D_2_O:H_2_O 4:1:0.2.

**Fig. S16**. A) time-kill profiles of allopeptimicin A against S. *aureus* ATCC6538P; B) Impact of allopeptimicins A on macromolecules synthesis by *S. aureus* cells: DNA (blue diamonds), RNA (orange squares), protein (grey triangles), and cell wall (yellow circles); C) growth curve of *S. aureus* ATCC6538P (dashed lines) and mutant R15.5^36^ (solid lines) in the presence of allopeptimicin A.

**Fig. S17**. Sensibility test of *Actinollomurus* ID145808 to allopeptimicins. Amount (in µg) of allopeptimicins A (panel A) and B (panel B) spotted in 10 µL is indicated on the plates. Bottom portion of each plate: Ap (0.1 µg apramycin in 10 µl) and 60% MeOH (10 µL) used as positive and negative controls, respectively.

## References

1. M. Miethke, M. Pieroni, T. Weber, M. Brönstrup, P. Hammann, L. Halby, P.B. Arimondo, P. Glaser, B. Aigle, H.B. Bode, R. Moreira, Y. Li, A. Luzhetskyy, M.H. Medema, J.L. Pernodet, M. Stadler, J.R. Tormo, O. Genilloud, A.W. Truman, K.J. Weissman, E. Takano, S. Sabatini, E. Stegmann, H. Brötz-Oesterhelt, W. Wohlleben, M. Seemann, M. Empting, A.K.H. Hirsch, B. Loretz, C.M. Lehr, A. Titz, J. Herrmann, T. Jaeger, S. Alt, T. Hesterkamp, M. Winterhalter, A. Schiefer, K. Pfarr, A. Hoerauf, H. Graz, M. Graz, M. Lindvall, S. Ramurthy, A. Karlén, M. van Dongen, H. Petkovic, A. Keller, F. Peyrane, S. Donadio, L. Fraisse, L.J.V. Piddock, I.H. Gilbert, H.E. Moser and R. Müller, Nat Rev Chem., 2021, 19, 1–24.

2. D.J. Newman and G.M. Cragg, J. Nat. Prod., 2020, 83, 770–803.

3. C. Kealey, C.A. Creaven, C.D. Murphy and C.B. Brady, Biotechnol. Lett., 2017, 39, 805–817.

4. M.H. Medema and M.A. Fischbach, Nat. Chem. Biol., 2015, 11, 639–648.

5. M. Wang, J.J. Carver, V.V. Phelan, L.M. Sanchez, N. Garg, Y. Peng, D.D. Nguyen, J. Watrous, C.A. Kapono, T. Luzzatto-Knaan, C. Porto, A. Bouslimani, A.V. Melnik, M.J. Meehan, W.T. Liu, M. Crüsemann, P.D. Boudreau, E. Esquenazi, M. Sandoval-Calderón, R.D. Kersten, L.A. Pace, R.A. Quinn, K.R. Duncan, C.C. Hsu, D.J. Floros, R.G. Gavilan, K. Kleigrewe, T. Northen, R.J. Dutton, D. Parrot, E.E. Carlson, B. Aigle, C.F. Michelsen, L. Jelsbak, C. Sohlenkamp, P. Pevzner, A. Edlund, J. McLean, J. Piel, B.T. Murphy, L. Gerwick, C.C. Liaw, Y.L. Yang, H.U. Humpf, M. Maansson, R.A. Keyzers, A.C. Sims, A.R. Johnson, A.M. Sidebottom, B.E. Sedio, A. Klitgaard, C.B. Larson, p CAB, D. Torres-Mendoza, D.J. Gonzalez, D.B. Silva, L.M. Marques, D.P. Demarque, E. Pociute, E.C. O’Neill, E. Briand, E.J.N. Helfrich, E.A. Granatosky, E. Glukhov, F. Ryffel, H. Houson, H. Mohimani, J.J. Kharbush, Y. Zeng, J.A. Vorholt, K.L. Kurita, P. Charusanti, K.L. McPhail, K.F. Nielsen, L. Vuong, M. Elfeki, M.F. Traxler, N. Engene, N. Koyama, O.B. Vining, R. Baric, R.R. Silva, S.J. Mascuch, S. Tomasi, S. Jenkins, V. Macherla, T. Hoffman, V. Agarwal, P.G. Williams, J. Dai, R. Neupane, J. Gurr, A.M.C. Rodríguez, A. Lamsa, C. Zhang, K. Dorrestein, B.M. Duggan, J. Almaliti, P.M. Allard, P. Phapale, L.F. Nothias, T. Alexandrov, M. Litaudon, J.L. Wolfender, J.E. Kyle, T.O. Metz, T. Peryea, D.T. Nguyen, D. VanLeer, P. Shinn, A. Jadhav, R. Müller, K.M. Waters, W. Shi, X. Liu, L. Zhang, R. Knight, P.R. Jensen, B.O. Palsson, K. Pogliano, R.G. Linington, M. Gutiérrez, N.P. Lopes, W.H. Gerwick, B.S. Moore, P.C. Dorrestein and N. Bandeira, Nat Biotechnol., 2016, 34, 828–837.

6. B.B. Misra. Metabolomics., 2021, 17, 49.

7. S.A. Jarmusch, J.J.J. van der Hooft, P.C. Dorrestein and A.K. Jarmusch, Nat Prod Rep., 2021, 38, 2066–2082.

8. M.M. Zdouc, M. Iorio, S.I. Maffioli, M. Crüsemann, S. Donadio and M. Sosio, J Nat Prod., 2021, 84, 204–219.

9. M.M. Zdouc, M.M. Alanjary, G.S. Zarazúa, S.I. Maffioli, M. Crüsemann, M.H. Medema, S. Donadio and M. Sosio, Cell Chem Biol., 2021, 28, 733-739.e4.

10. J.J. Hug, N.A. Frank, C. Walt, P. Šenica, F. Panter and R. Müller, Molecules., 2021, 26, 7483.

11. M. Iorio, A. Tocchetti, J.C. Cruz, G. Del Gatto, C. Brunati, S.I. Maffioli, M. Sosio and S. Donadio, Antibiotics (Basel). 2018, 7, 47,

12. M. Iorio, J.C. Cruz, M. Simone, A. Bernasconi, C. Brunati, M. Sosio, S. Donadio and S.I. Maffioli, J Nat Prod., 2017, 80, 819–827.

13. J.C. Cruz, S.I. Maffioli, A. Bernasconi, C. Brunati, E. Gaspari, M. Sosio, E. Wellington and S. Donadio, J Antibiot (Tokyo)., 2017, 70, 73–78.

14. J.C. Cruz, M. Iorio, P. Monciardini, M. Simone, C. Brunati, E. Gaspari, S.I. Maffioli, E. Wellington, M. Sosio and S. Donadio, J Nat Prod., 2015, 78, 2642–7.

15. C. Mazzetti, M. Ornaghi, E. Gaspari, S. Parapini, S.I. Maffioli, M. Sosio and S. Donadio, J Nat Prod. 2012, 75, 1044–50.

16. R. Pozzi, M. Simone, C. Mazzetti, S.I. Maffioli, P. Monciardini, L. Cavaletti, R. Bamonte, M. Sosio and S. Donadio, S. J Antibiot (Tokyo)., 2011, 64, 133–9.

17. H. Bruns, M. Crüsemann, A.C. Letzel, M. Alanjary, J.O. McInerney, P.R. Jensen, S. Schulz, B.S. Moore and N. Ziemert, ISME J. 2018, 12, 320–329.

18. A. Miyanaga, F. Kudo and T. Eguchi, Curr Opin Chem Biol., 2016, 35, 58–64.

19. F. Kudo, A. Miyanaga and T. Eguchi, Nat Prod Rep., 2014, 31, 1056–73.

20. M. Takaishi, F. Kudo and T. Eguchi, J Antibiot, 2013, 66, 691–699.

21. J. Cieślak, A. Miyanaga, R. Takaku, M. Takaishi, K. Amagai, F. Kudo and T. Eguchi, T. Proteins, 2017, 85, 1238–1247.

22. Z. Yin and J.S. Dickshat, Nat. Prod. Rep., 2021, 38, 1445–1468.

23. F. Sun, S. Xu, F. Jiang and W. Liu, Appl Microbiol Biotechnol., 2018, 102, 2225–2234.

24. S.R. Lee, H. Guo, J.S. Yu, M. Park, H. Dahse, W.H. Jung, C. Beemelmanns and K.H. Kim, Org. Chem. Front., 2021,8, 4791–4798.

25. J. Ma, Z. Wang, H. Huang, M. Luo, D. Zuo, B. Wang, A. Sun, Y.Q. Cheng, C. Zhang and J. Ju, Angew Chem Int Ed Engl., 2011, 50, 7797–802.

26. D.G. Fujimori, S. Hrvatin, C.S. Neumann, M. Strieker, M.A. Marahiel and C.T. Walsh Ct, Proc Natl Acad Sci U S A. 2007, 104,16498–503.

27. C.T. Walsh, R.V. O’Brien and C. Khosla, Angewandte Chemie International Edition., 2013, 52, 7098–7124.

28. Q. Li, X. Qin, J. Liu, C. Gui, B. Wang, J. Li and J. Ju, J Am Chem Soc., 2016, 138, 408–15.

29. K.D. Morgan, R.J. Andersen and K.S. Ryan Ks. Nat Prod Rep., 2019, 36, 1628–1653.

30. H. Zhao, L. Wang, D. Wan, J. Qi, R. Gong, Z. Deng and W. Chen, W. Microbial cell factories., 2016, 15, 160.

31. R. Li, E.P. Lloyd, K.A. Moshos and C.A. Townsend, ChemBioChe., 2014, 15, 320 – 331.

32. Y. Kato, T. Suzuki, T. Ida and K. Maebashi, Journal of Antimicrobial Chemotherapy., 2010, 65, 37–45.

33. B.R. Miller, E.J. Drake, C. Shi, C.C. Aldrich and A.M. Gulick, Journal of Biological Chemistry, 2016, 43, 22559–22571.

34. S.C. Curran, J.H. Pereira, M.J. Baluyot, J. Lake, H. Puetz, D.J. Rosenburg, P. Adams and J.D. Keasling, Biochemistry., 2020, 59, 1630–1639.

35. S.A. Kautsar, K. Blin, S. Shaw, J.C. Navarro-Muñoz, B.R. Terlouw, J.J.J. van der Hooft, J.A. van Santen, V. Tracanna, H.G. Suarez Duran, V. Pascal Andreu, N. Selem-Mojica, M. Alanjary, S.L. Robinson, G. Lund, S.C. Epstein, A.C. Sisto, L.K. Charkoudian, J. Collemare, R.G. Linington, T. Weber and M.H. Medema Mh, Nucleic Acids Res., 2020, 48, D454–D458.

36. A. Tocchetti, M. Iorio, Z. Hamid, A. Armirotti, A. Reggiani and S. Donadio, Molecules., 2021, 26, 6764.

37. J. Jokela, L. Oftedal, L. Herfindal, P. Permi, M. Wahlsten, S.O. Døskeland and K. Sivonen, PLoS One., 2012, 7, e41222.

38. K. Petříčková, S. Pospíšil, M. Kuzma, T. Tylová, M. Jágr, P. Tomek, A. Chroňáková, E. Brabcová, L. Anděra, V. Krištůfek and M. Petříček, Chembiochem., 2014, 15, 1334–45.

39. B. Kunze, H. Reichenbach, R. Müller and G. Höfle, J Antibiot (Tokyo)., 2005, 58, 244–51.

40. J. Niggemann, N. Bedorf, U. Flörke, H. Steinmetz, K. Gerth, H. Reichenbach and G. Höfle, EurJOC, 2005, 23, 5013–5018.

41. M.A. Exposito, B. Lopez, R. Fernandez, M.J. Vazquez, C. Debitus, T. Iglesias, C. Jimenez, E. Quinoa and R. Riguera, Tetrahedron., 1998, 54, 7539–7550.

42. E.W. Schmidt, C. Raventos-Suarez, M. Bifano, A.T. Menendez, C.R. Fairchild and D.J. Faulkner, J Nat Prod., 2004, 67, 475–8.

43. Y. Okamoto, M. Ojika, S. Kato and Y. Sakagami, Tetrahedron., 2000, 56, 5813–5818.

44. B. Seger, S. Sturm, H. Stuppner, T.M. Butt and H. Strasser, J Chromatogr A., 2004, 1061, 35–43.

45. M.H. Medema, T. de Rond and B.S. Moore, Nat Rev Genet., 2021, 22, 553–571.

46. A.L. Lukowski, N. Denomme, M.E. Hinze, S. Hall, L.L. Isom, and A. Narayan, ACS Chemical Biology., 2019, 14, 941–948.

47. S. Donadio, P. Monciardini and M. Sosio M, Methods Enzymol. 2009, 458, 3–28.

48. T. Pluskal, S. Castillo, A. Villar-Briones and M. Oresic, BMC Bioinformatics., 2010, 23, 11:395.

49. P. Shannon, A. Markiel, O. Ozier, N.S. Baliga, J.T. Wang, D. Ramage, N. Amin, B. Schwikowski and T. Ideker, Genome Res., 2003, 13, 2498–504.

50. R. Piccolomini, G. Di Bonaventura, G. Catamo, F. Carbone and M. Neri, Journal of Clinical Microbiology., 1997, 35, 1842–1846.

51. S.I. Maffioli, P. Monciardini, B. Catacchio, C. Mazzetti, D. Münch, C. Brunati, H.G. Sahl and S. Donadio S. ACS Chem Biol., 2015, 10, 1034–42.

52. P. Monciardini, A. Bernasconi, M. Iorio, C. Brunati, M. Sosio, L. Campochiaro, P. Landini, S.I. Maffioli and S. Donadio, J Nat Prod., 2019, 82, 35–44.

53. K. Blin, S. Shaw, A.M. Kloosterman, Z. Charlop-Powers, G.P. van Wezel, M.H. Medema and T. Weber, Nucleic Acids Res., 2021, 49, W29–W35.

54. P. Mazza, P. Monciardini, L. Cavaletti, M. Sosio and S. Donadio, Microb Ecol., 2003, 45, 362–72.

