## Supplemental information for "Allopeptimicins: unique antibacterial metabolites generated by hybrid PKS-NRPS, with original self-defense mechanism in *Actinoallomurus*"

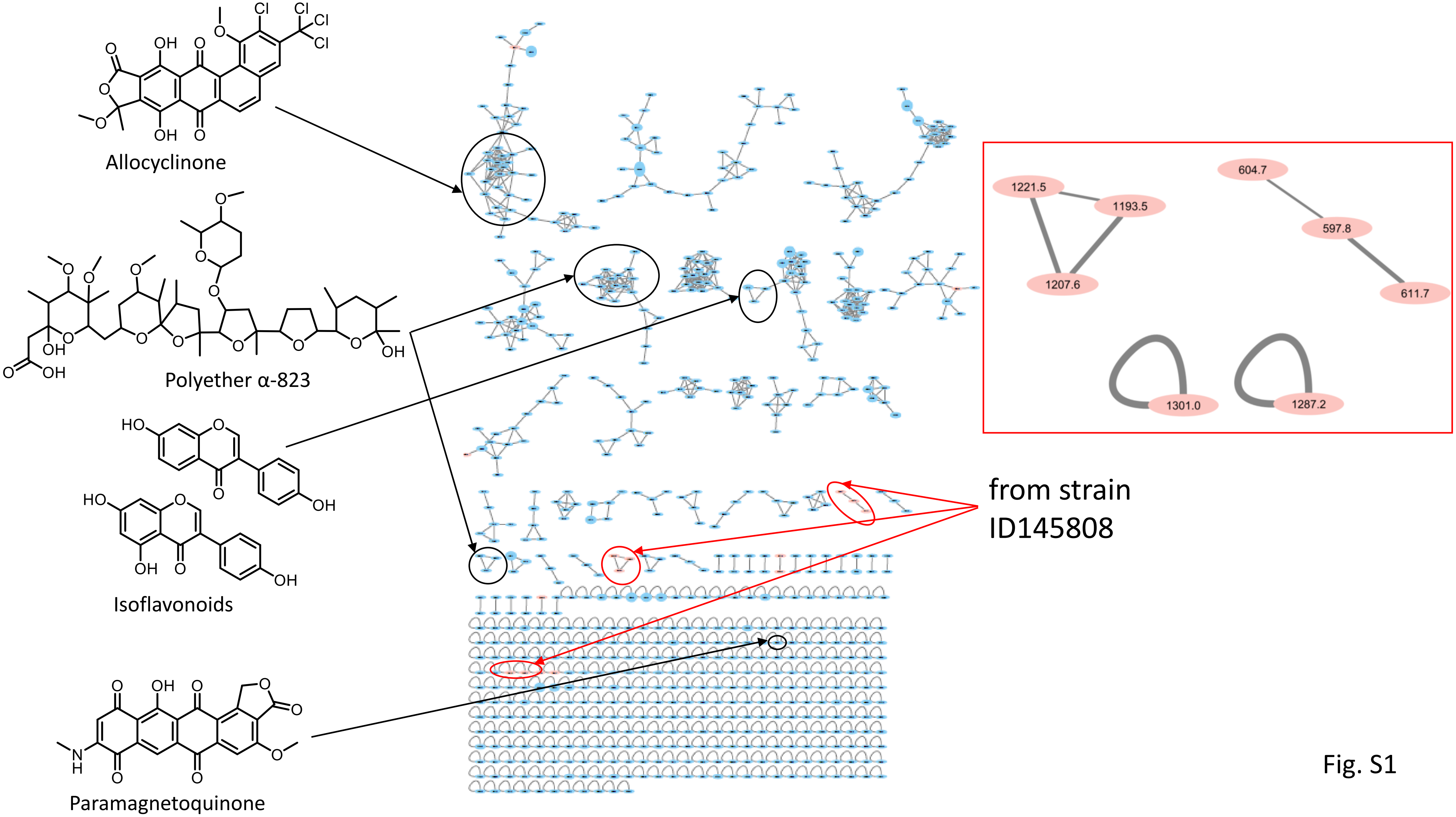

Allocyclinone
Polyether α-823
Isoflavonoids
Paramagnetoquinone
from strain ID145808
1221.5
1193.5
1207.6
604.7
597.8
611.7
1301.0
1287.2


Fig. S1

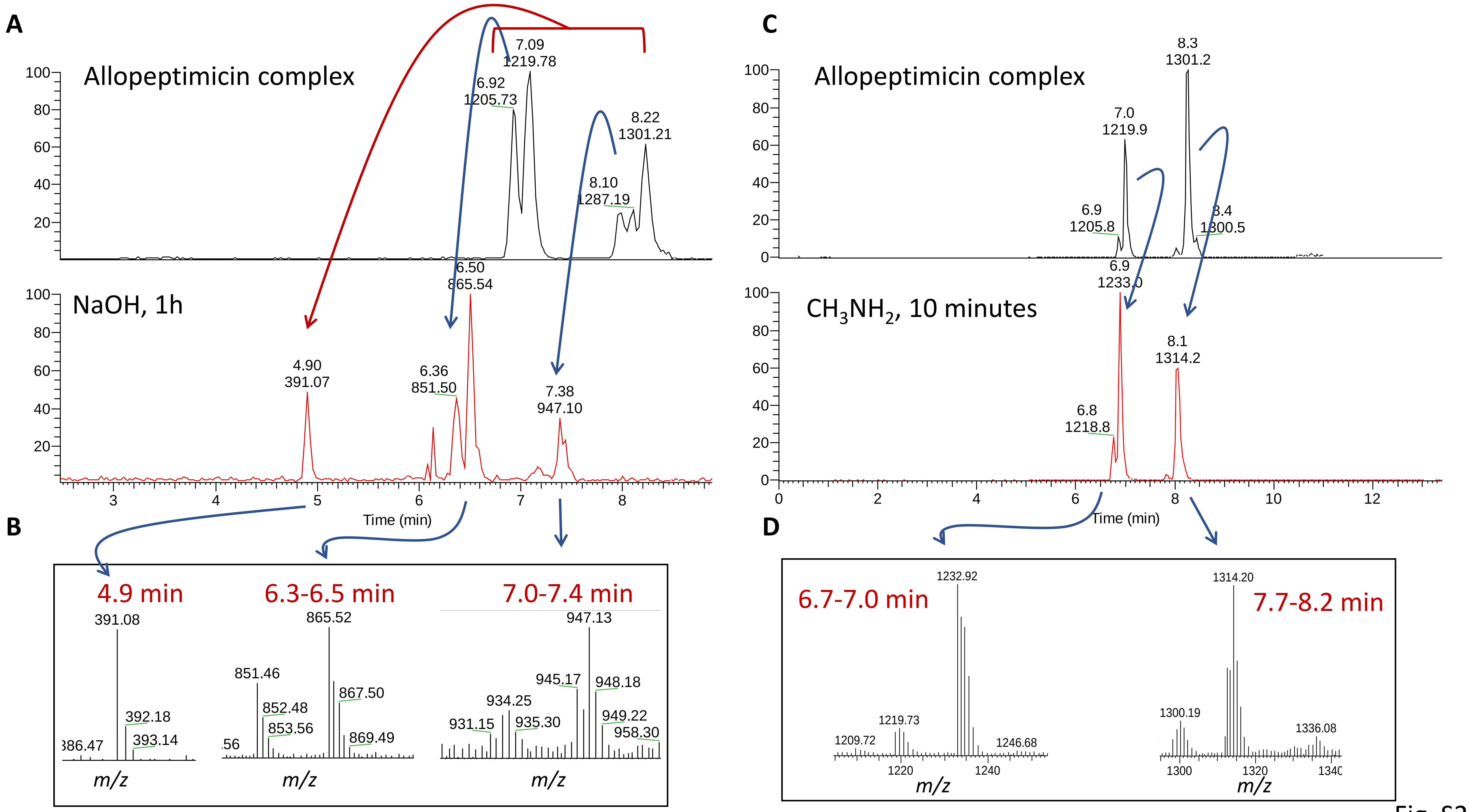
A
Allopeptimicin complex
7.09
1219.78
6.92
1205.73
8.22
1301.21
8.10
1287.19
NaOH, 1h
6.50
865.54
4.90
391.07
6.36
851.50
7.38
947.10
Time (min)
B
4.9 min
391.08
392.18
393.14
6.3-6.5 min
865.52
851.46
852.48
853.56
867.50
869.49
7.0-7.4 min
947.13
945.17
948.18
934.25
931.15
935.30
949.22
958.30
m/z
C
Allopeptimicin complex
8.3
1301.2
7.0
1219.9
6.9
1205.8
1300.5
CH3NH2, 10 minutes
6.9
8.1
1314.2
6.8
1218.8
Time (min)
D
6.7-7.0 min
1232.92
1219.73
1209.72
1246.68
7.7-8.2 min
1314.20
1300.19
1336.08
m/z

Fig. S2

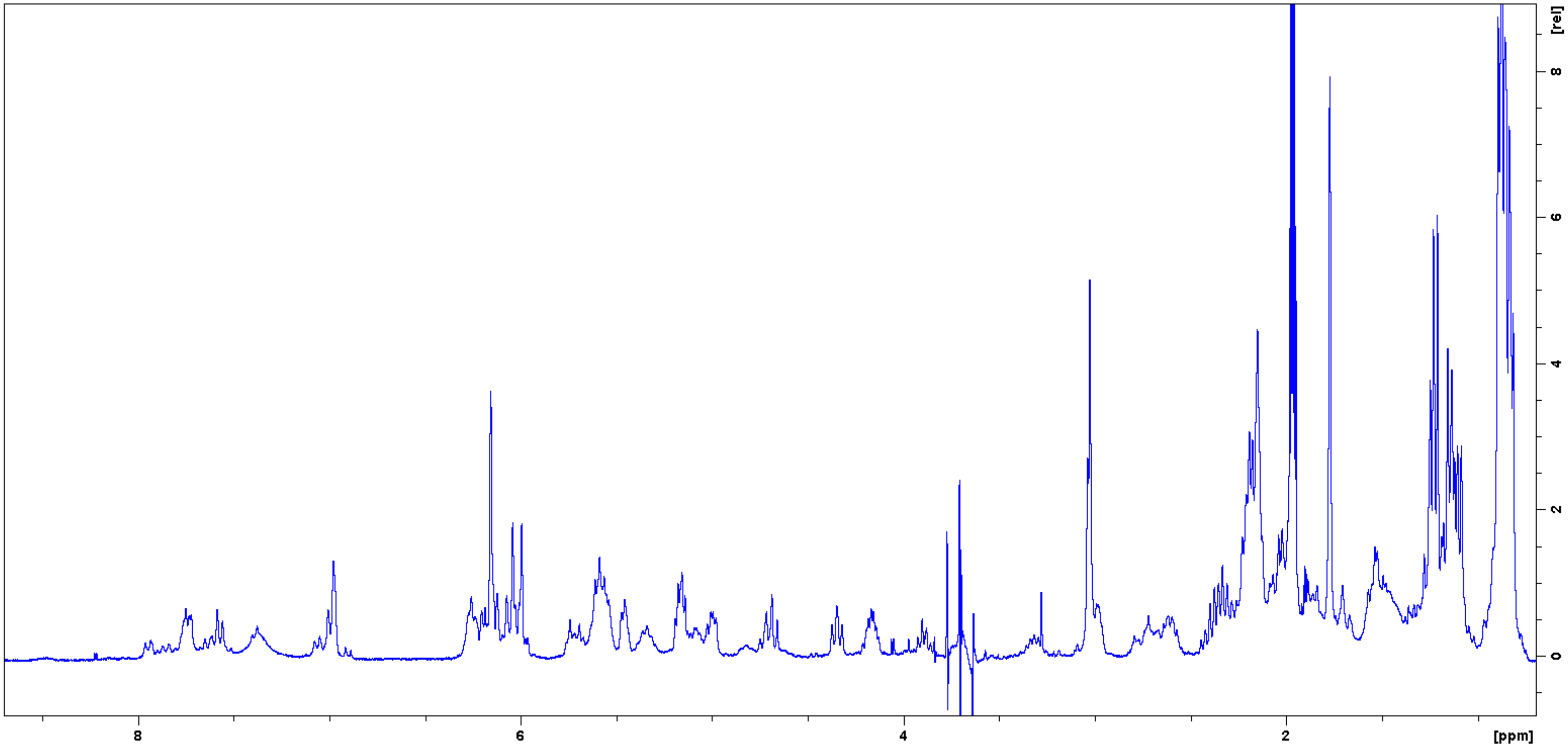


Fig. S3

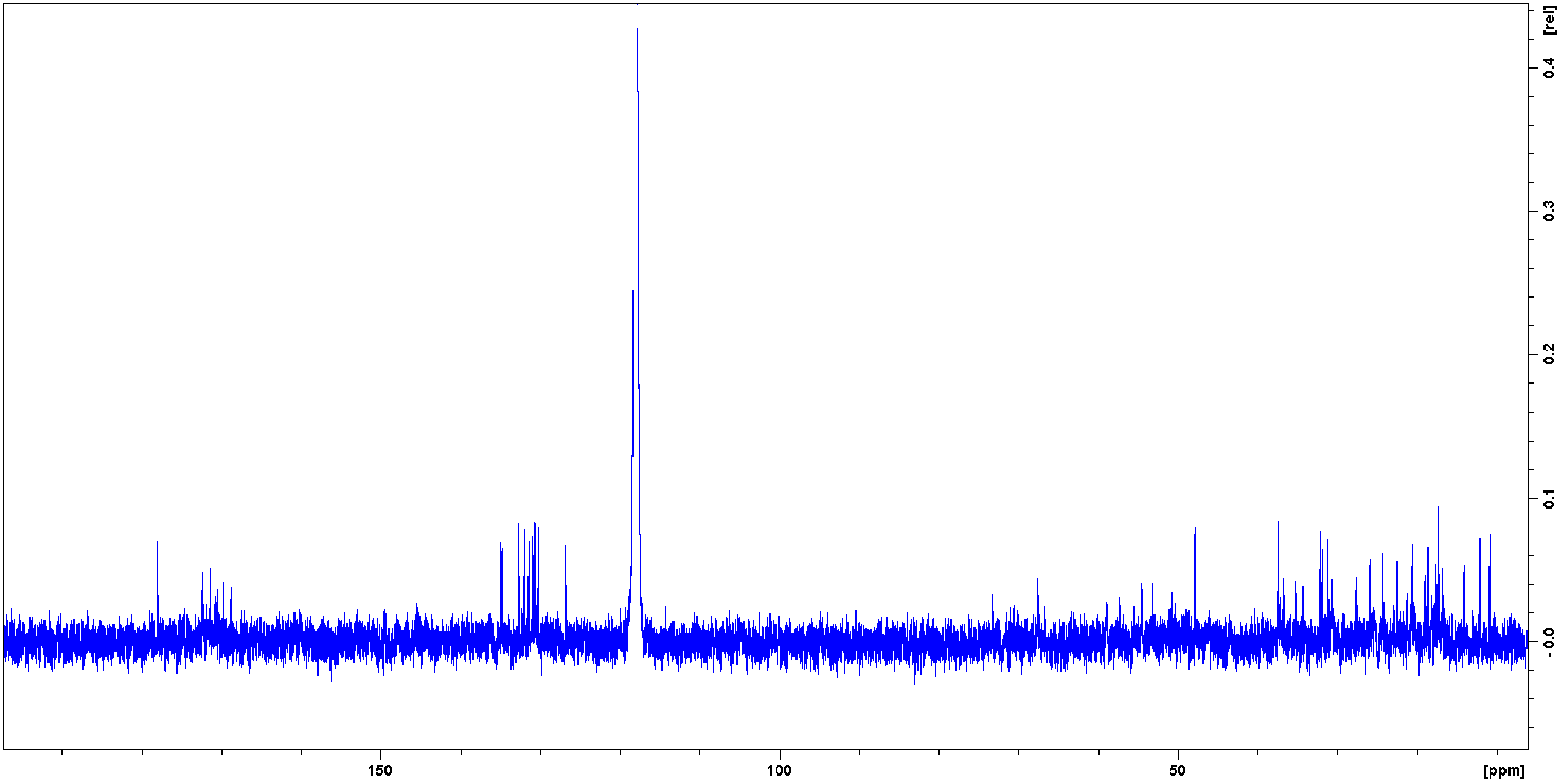


Fig. S4

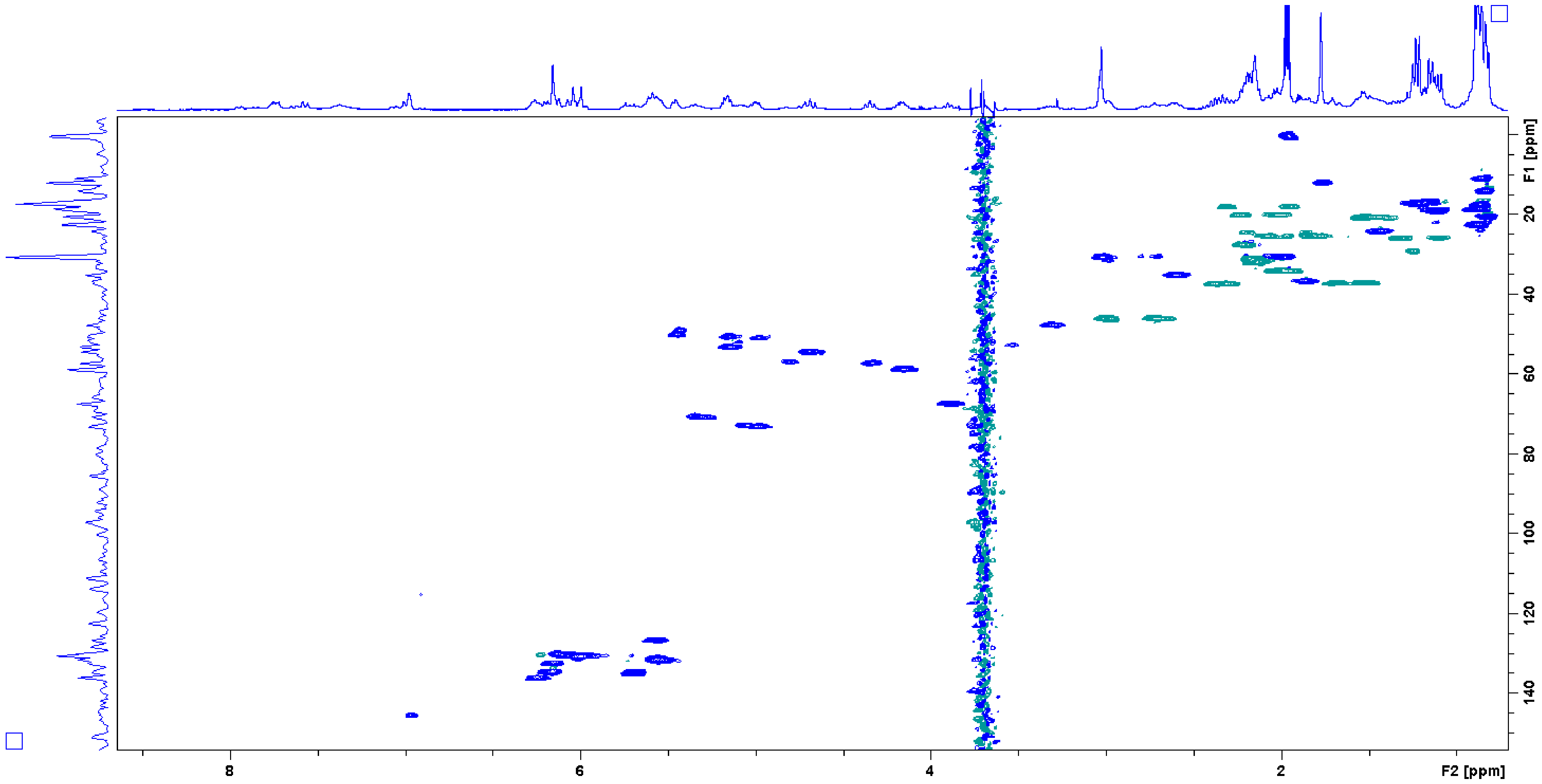


Fig. S5

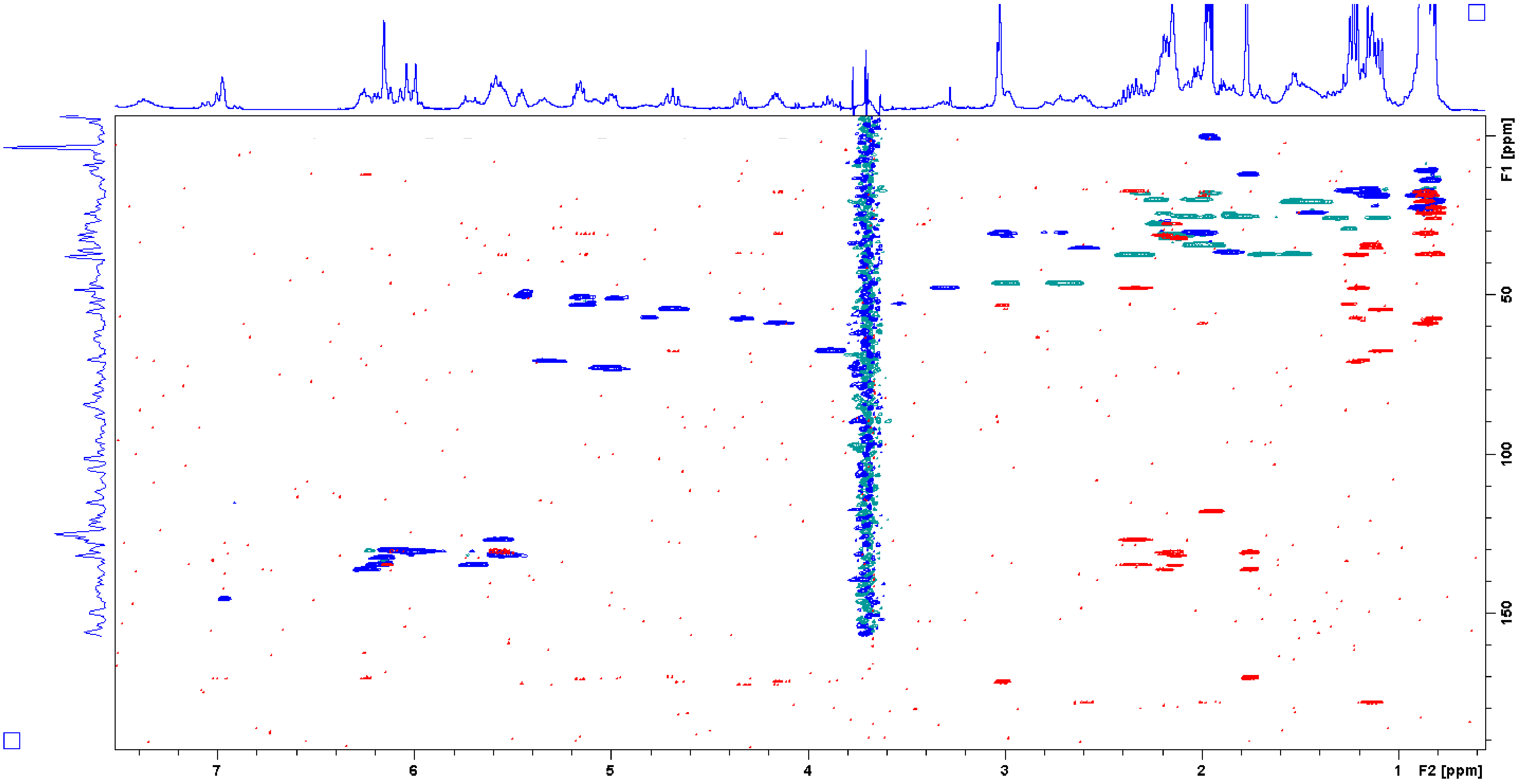


Fig. S6

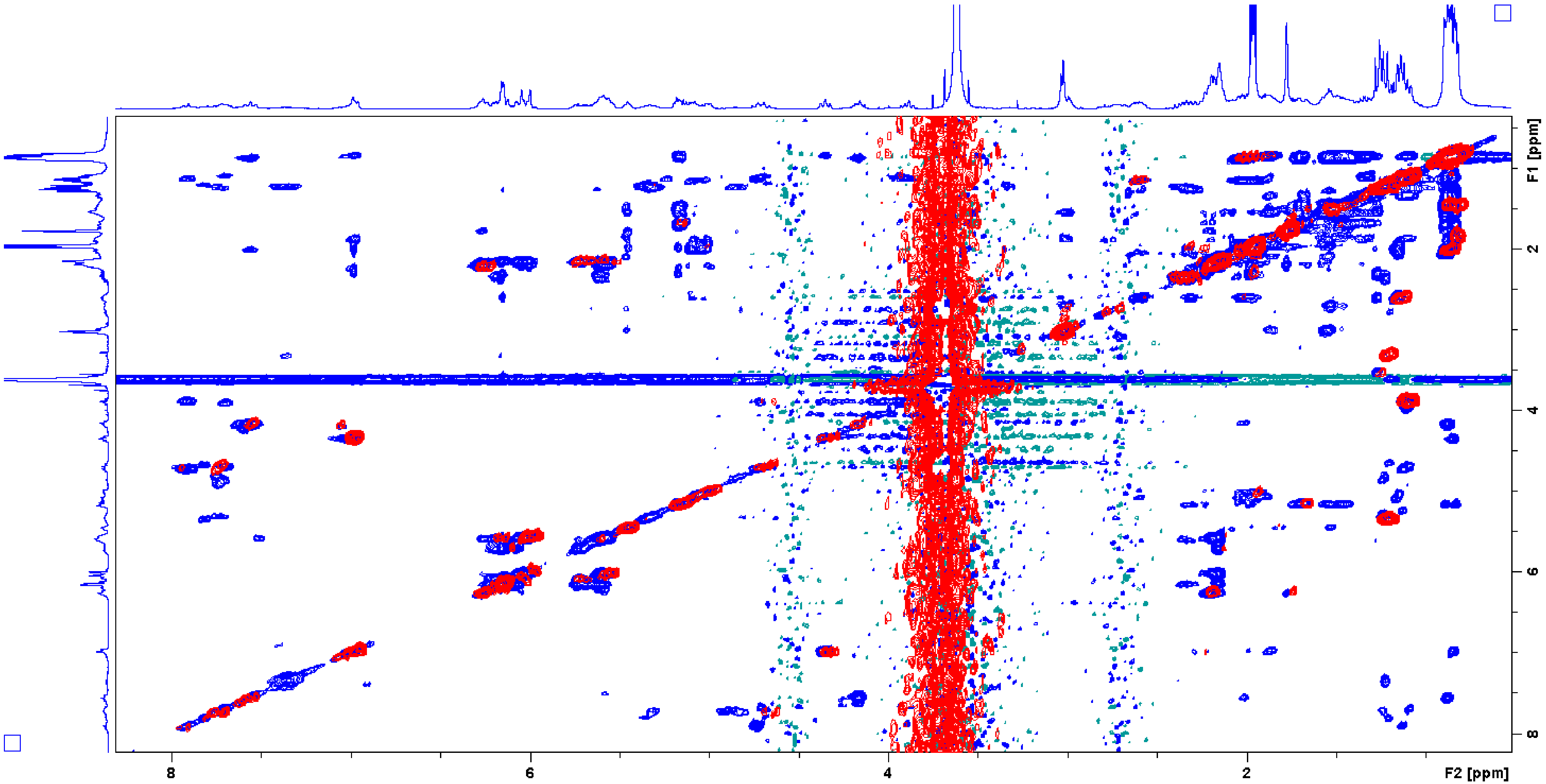


Fig. S7

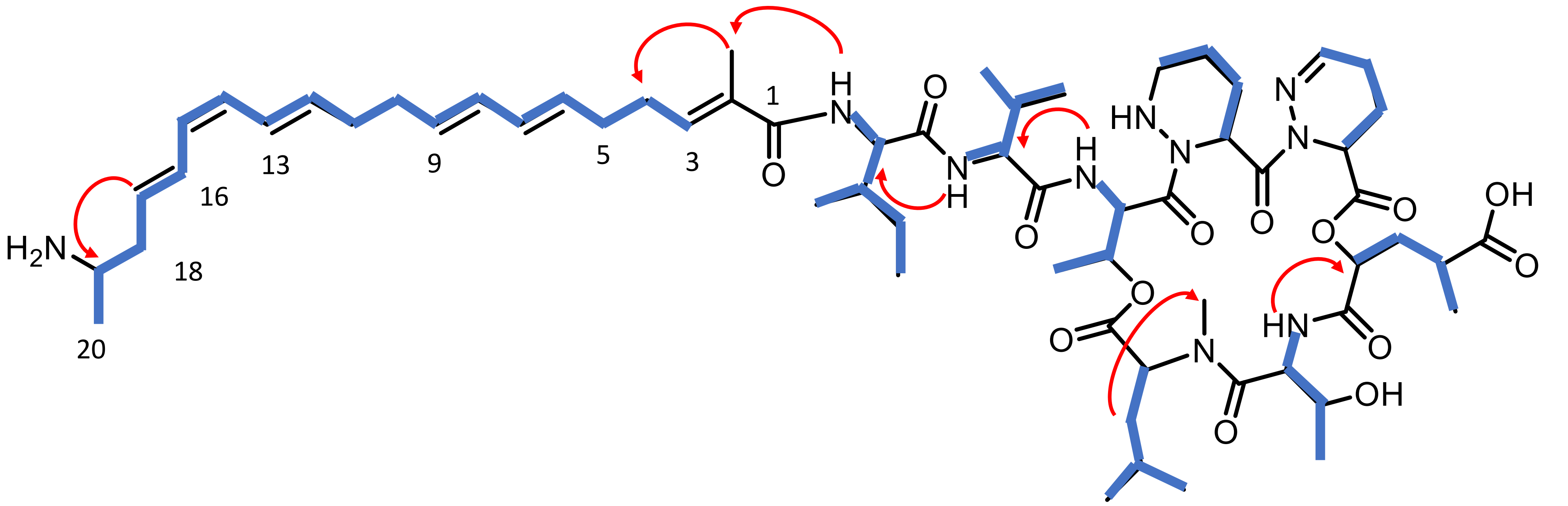


Fig. S8

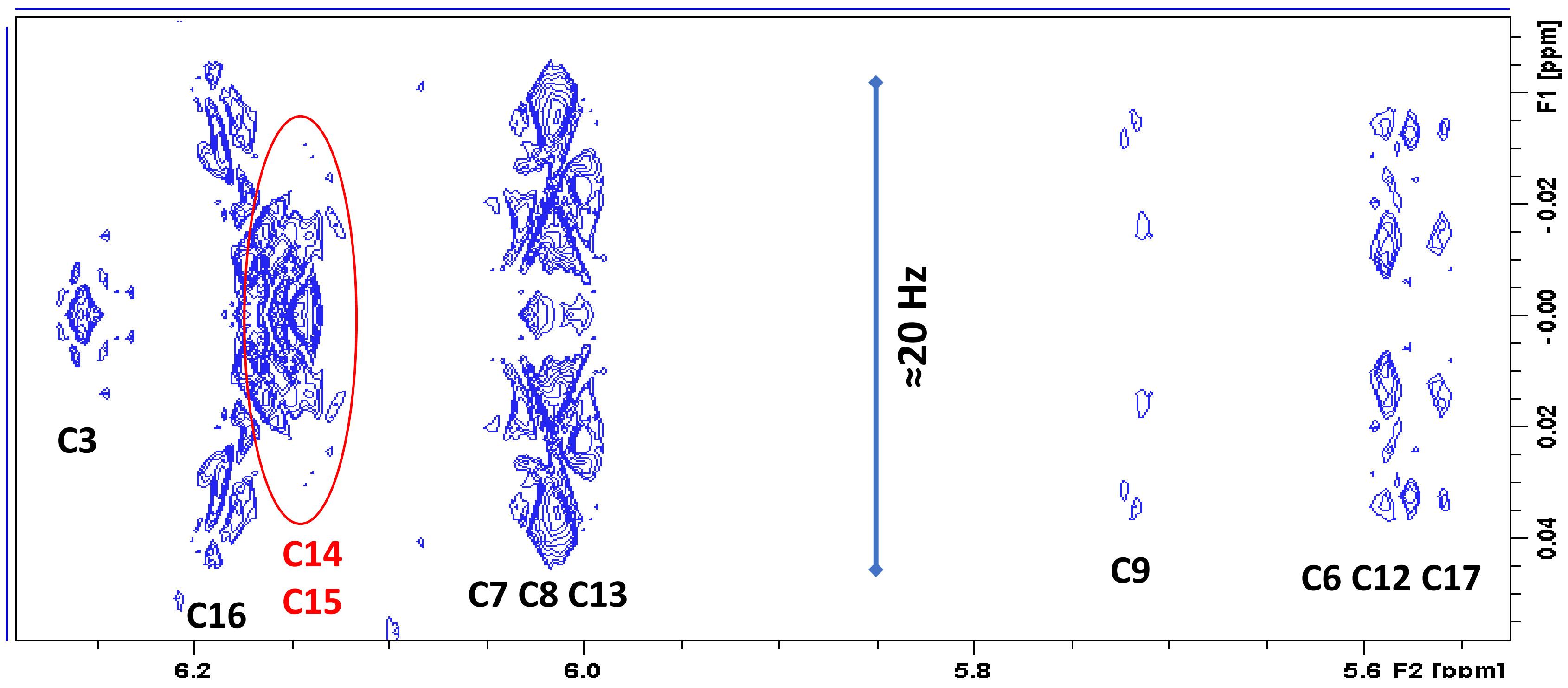
C3
C16
C14
C15
C7 C8 C13
≈20 Hz
C9
C6 C12 C17
6.2
6.0
5.8
5.6 F2 [ppm]
F1 [ppm]
-0.02
-0.00
0.02
0.04

Fig. S9

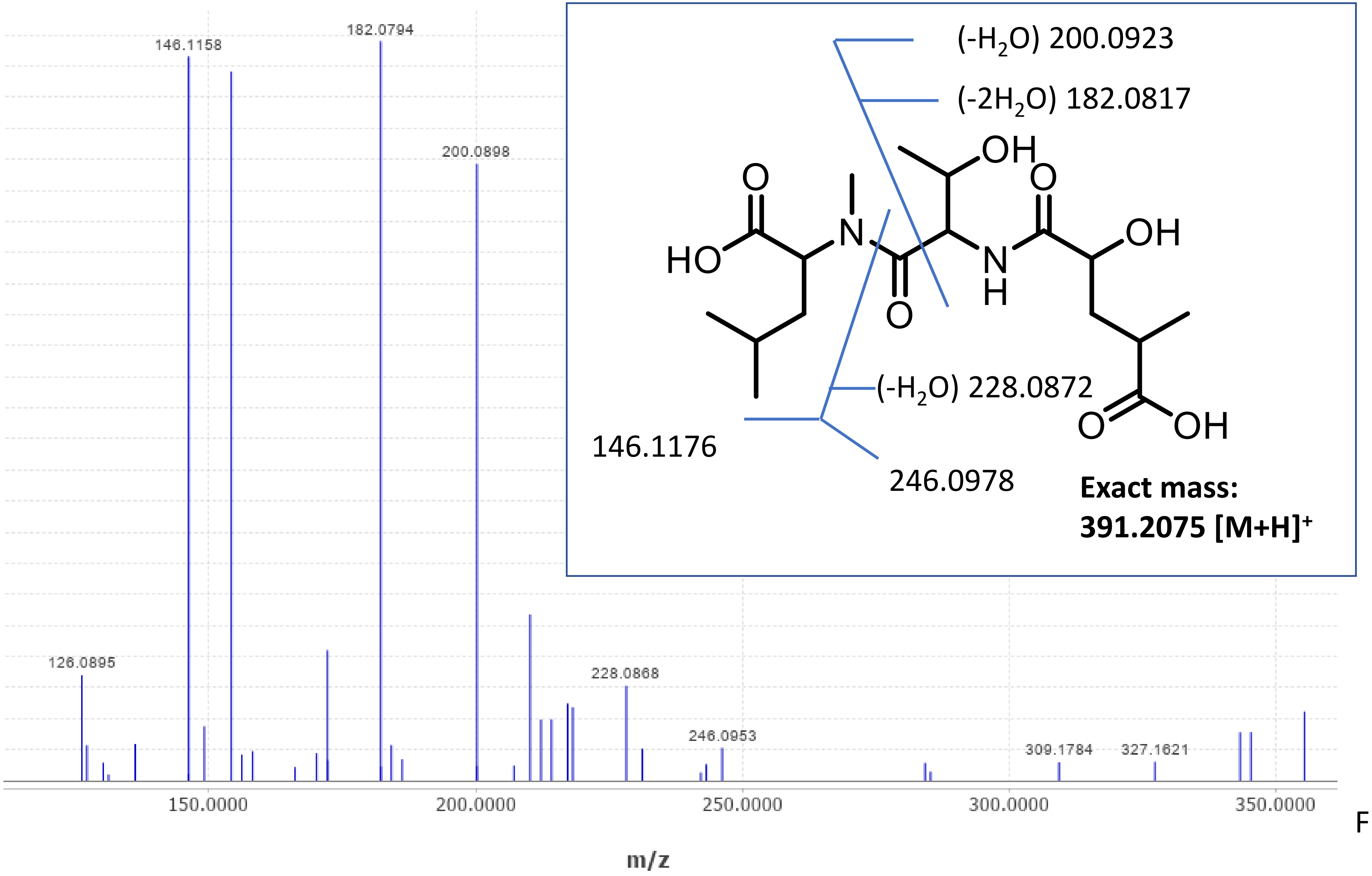

146.1158
182.0794
200.0898
126.0895
228.0868
246.0953
309.1784
327.1621
150.0000
200.0000
250.0000
300.0000
350.0000
m/z
(-H2O) 200.0923
(-2H2O) 182.0817
(-H2O) 228.0872
146.1176
246.0978
Exact mass:
391.2075 [M+H]+


Fig. S10

A

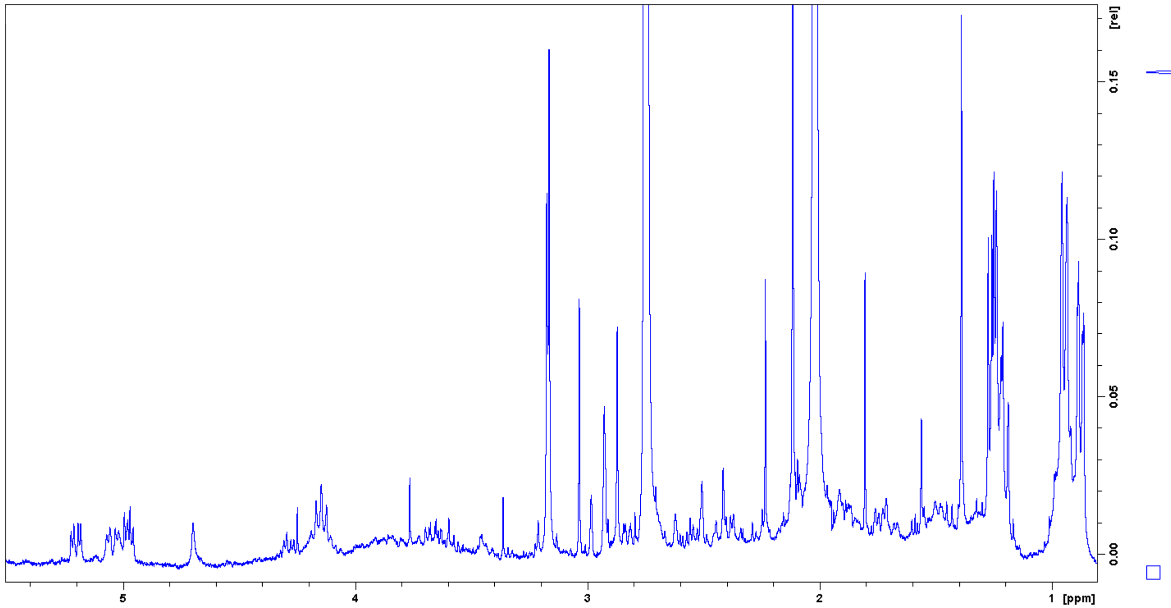

B

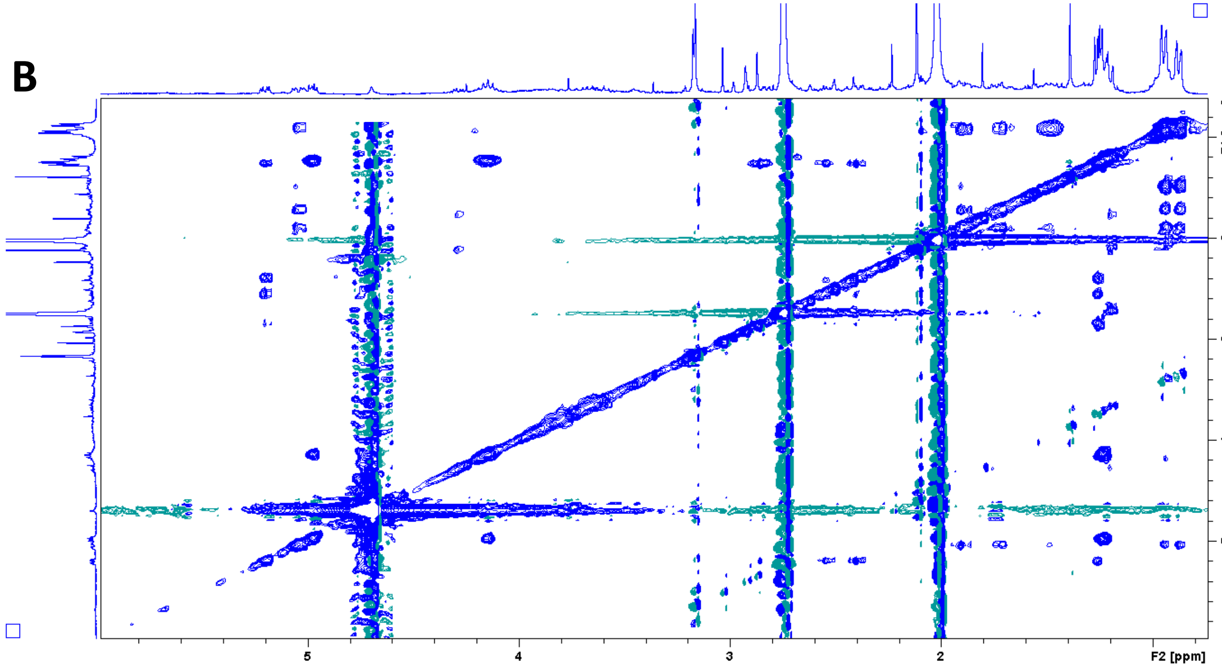


C

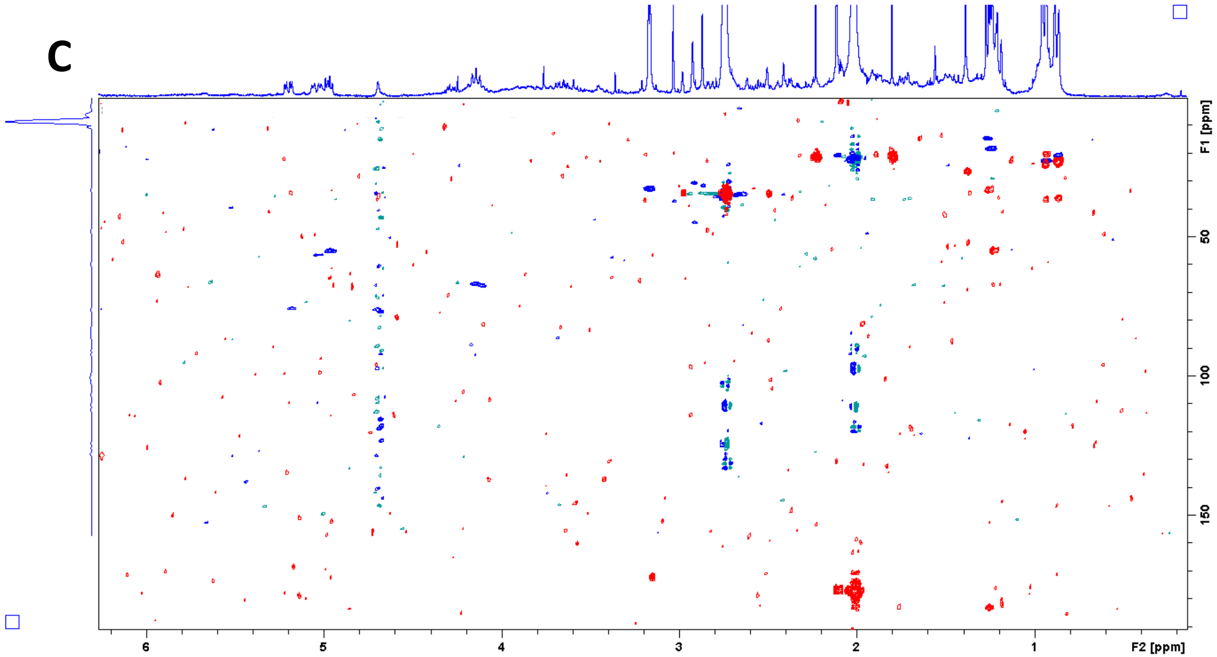


D

| unit | position | Hydrolytic fragment δH | δC |
|---|---|---|---|
| Hmg_6 | 1 | | 168.4 |
| | 2 | 5.19 | 75.0 |
| | 3a | 2.33 | 33.7 |
| | 3b | 2.41 | |
| | 4 | 2.84 | 34.6 |
| | 5 | | 183.3 |
| | 4-Me | 1.26 | 14.6 |
| Thr_7 | NH | 7.31 | |
| | 1 | | 172.2 |
| | 2 | 4.98 | 55.3 |
| | 3 | 4.13 | 67.1 |
| | 4 | 1.23 | 18.6 |
| N-Me-Leu_8 | 1 | | 172.1 |
| | 2 | 5.03 | 56.3 |
| | 3a | 1.76 | 36.5 |
| | 3b | 1.92 | |
| | 4 | 1.48 | 24.4 |
| | 5 | 0.94 | 22.4 |
| | 6 | 0.87 | 20.5 |
| | N-Me | 3.17 | 32.4 |

Fig. S11

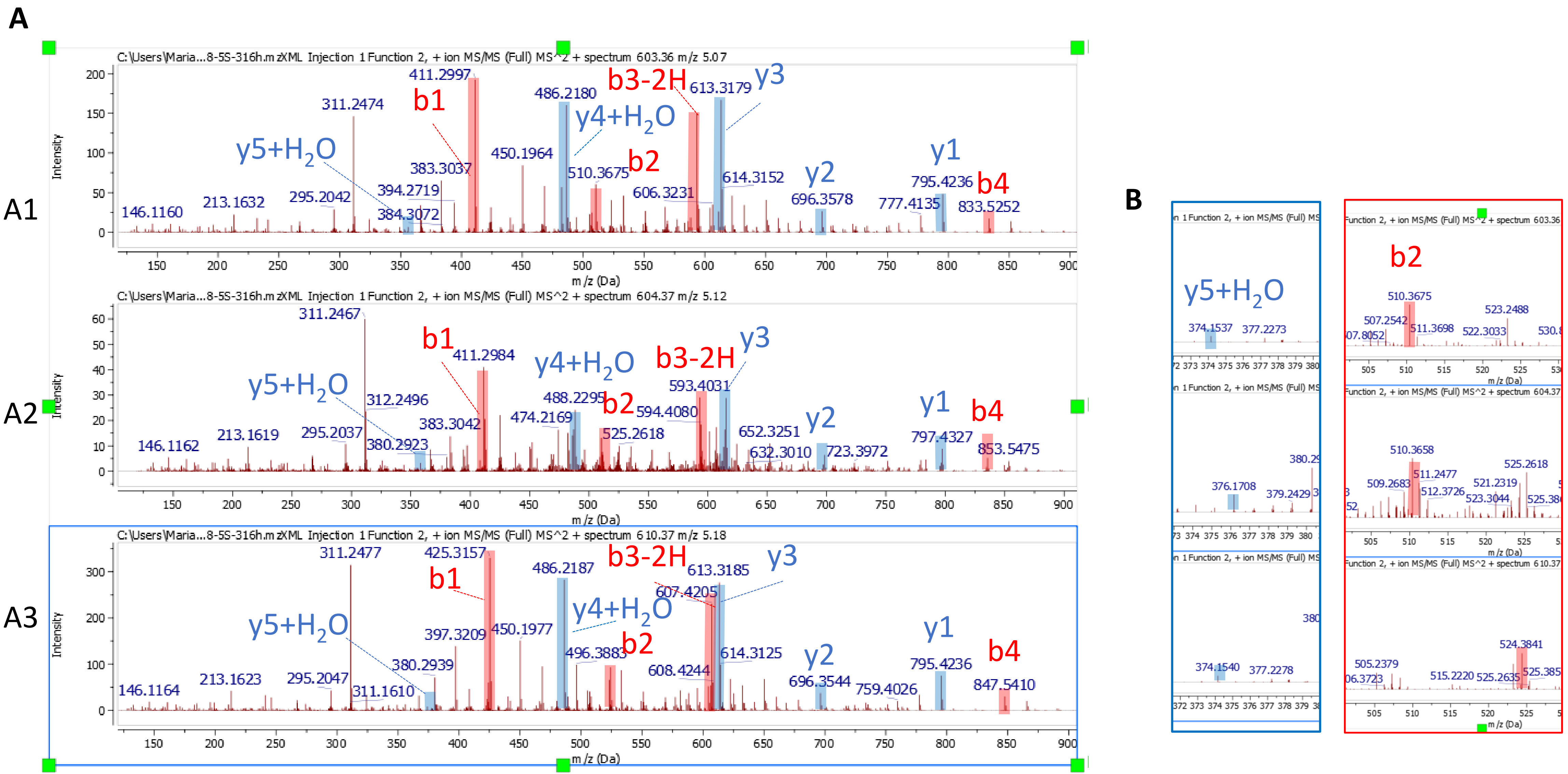
A
A1
A2
A3
B
C:\Users\Maria...8-5S-316h.mzXML Injection 1 Function 2, + ion MS/MS (Full) MS^2 + spectrum 603.36 m/z 5.07
C:\Users\Maria...8-5S-316h.mzXML Injection 1 Function 2, + ion MS/MS (Full) MS^2 + spectrum 604.37 m/z 5.12
C:\Users\Maria...8-5S-316h.mzXML Injection 1 Function 2, + ion MS/MS (Full) MS^2 + spectrum 610.37 m/z 5.18
Intensity
m/z (Da)
b1
b2
b3-2H
b4
y1
y2
y3
y4+H2O
y5+H2O
146.1160
213.1632
295.2042
311.2474
383.3037
384.3072
394.2719
411.2997
450.1964
486.2180
510.3675
606.3231
613.3179
614.3152
696.3578
777.4135
795.4236
833.5252
146.1162
213.1619
295.2037
311.2467
312.2496
380.2923
383.3042
411.2984
474.2169
488.2295
525.2618
593.4031
594.4080
632.3010
652.3251
723.3972
797.4327
853.5475
146.1164
213.1623
295.2047
311.1610
311.2477
380.2939
397.3209
425.3157
450.1977
486.2187
496.3883
607.4205
608.4244
613.3185
614.3125
696.3544
759.4026
795.4236
847.5410
374.1537
377.2273
376.1708
379.2429
374.1540
377.2278
507.2542
510.3675
511.3698
522.3033
523.2488
509.2683
510.3658
511.2477
512.3726
521.2319
523.3044
525.2618
505.2379
506.3723
515.2220
524.3841
525.2635

Fig. S12

**A**

| | Allopeptimicin A1 | | Allopeptimicin A2 | | Allopeptimicin A3 | |
|---|---|---|---|---|---|---|
| | found | calculated | found | calculated | found | calculated |
| $[M+H]^+$ | 1205.718 | 1205.718 | 1207.7348 | 1207.734 | 1219.734 | 1219.7337 |
| b4 | 833.5252 | 833.5284 | 835.538 | 835.544 | 847.541 | 847.5446 |
| y1 | 795.4236 | 795.4247 | 797.4327 | 797.4403 | 795.4236 | 795.4247 |
| y2 | 696.3578 | 696.3563 | 698.3643 | 698.3719 | 696.3544 | 696.3563 |
| y3 | 613.3179 | 613.3192 | 615.3274 | 615.3348 | 613.3185 | 613.3192 |
| b3-2H | 593.4056 | 593.4061 | 593.4031 | 593.4061 | 607.4205 | 607.4223 |
| b2 | 510.3675 | 510.3696 | 510.3658 | 510.3696 | 524.3841 | 524.3852 |
| y4 + $H_2O$ | 486.218 | 486.2195 | 488.2295 | 488.2351 | 486.2187 | 486.2195 |
| b1 | 411.2997 | 411.3012 | 411.2984 | 411.3012 | 425.3157 | 425.3168 |
| y5 + $H_2O$ | 374.1537 | 374.1558 | 376.1708 | 376.1714 | 374.154 | 374.1558 |

**B**

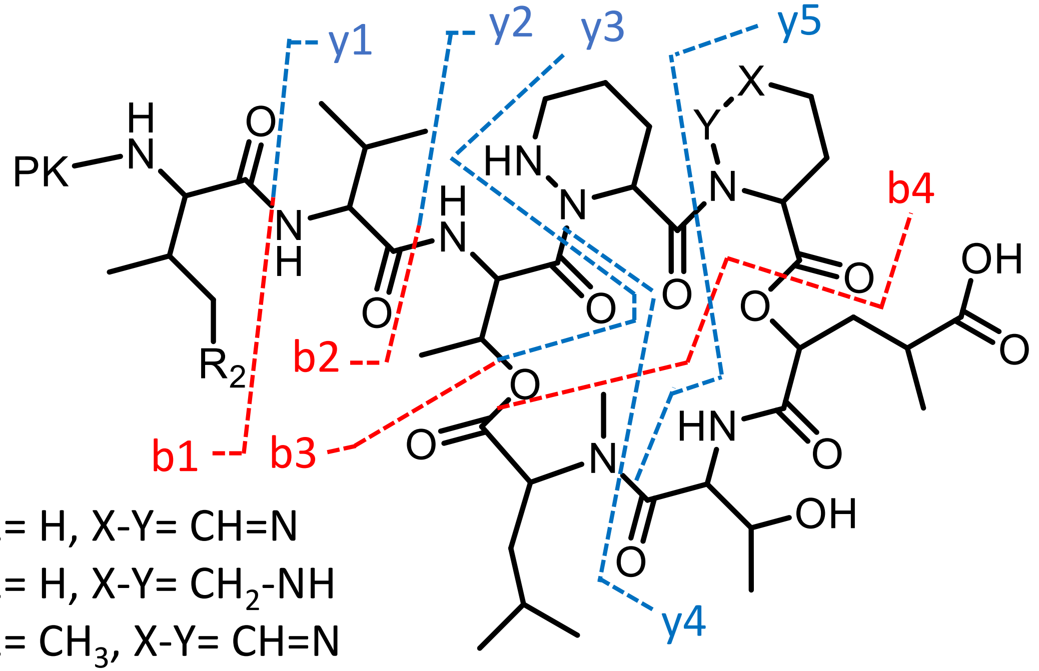


**A1**: R2= H, X-Y= CH=N

**A2**: R2= H, X-Y= $CH_2$-NH

**A3**: R2= $CH_3$, X-Y= CH=N

Fig. S13

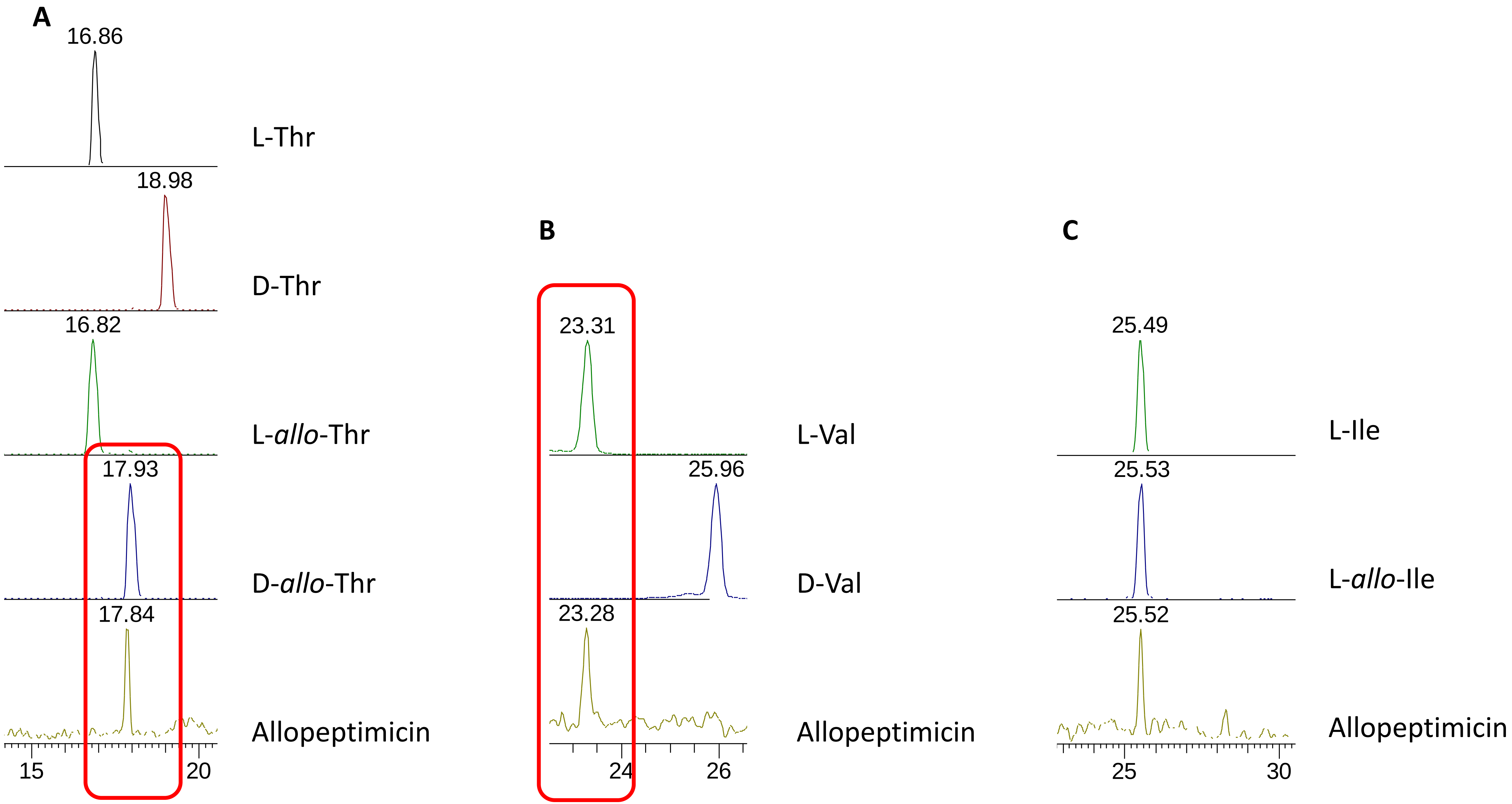
A
16.86
L-Thr
18.98
D-Thr
16.82
L-allo-Thr
17.93
D-allo-Thr
17.84
Allopeptimicin
15
20
B
23.31
L-Val
25.96
D-Val
23.28
Allopeptimicin
24
26
C
25.49
L-Ile
25.53
L-allo-Ile
25.52
Allopeptimicin
25
30

Fig. S14

A

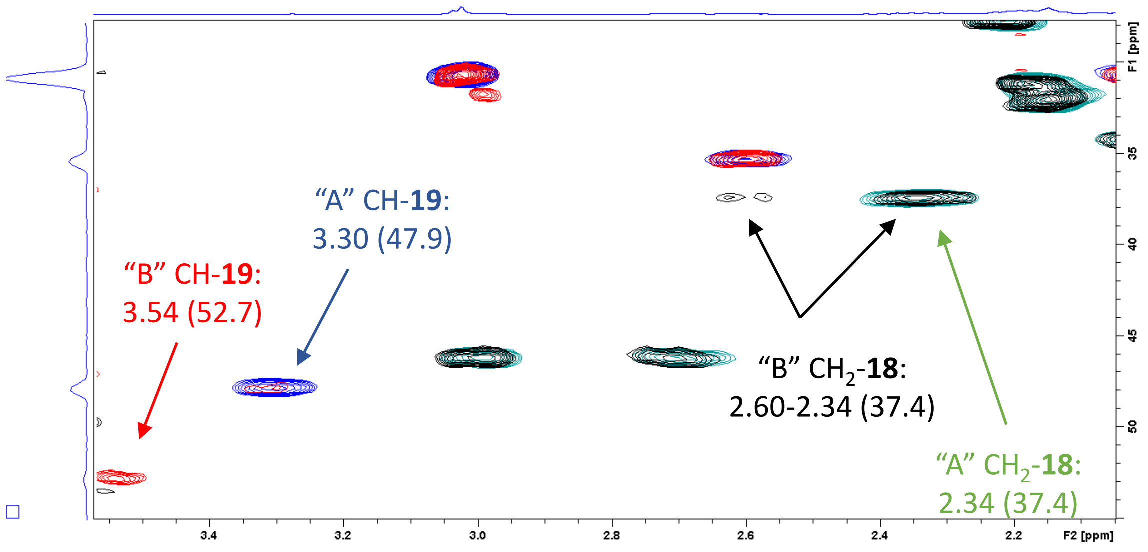
"A" CH-19:
3.30 (47.9)
"B" CH-19:
3.54 (52.7)
"B" CH2-18:
2.60-2.34 (37.4)
"A" CH2-18:
2.34 (37.4)
F1 [ppm]
F2 [ppm]

B

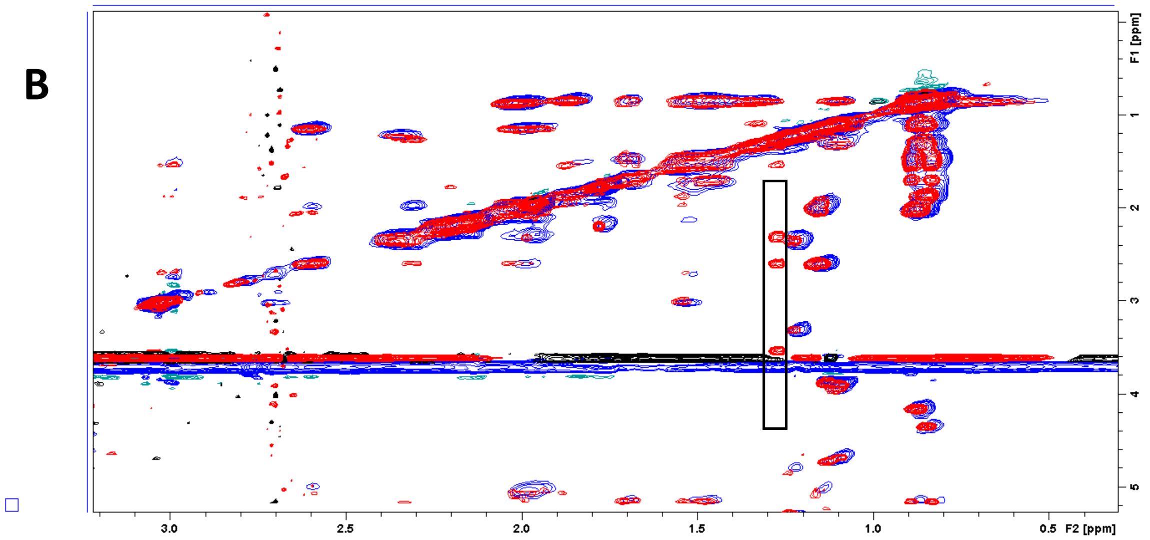
F1 [ppm]
F2 [ppm]

Fig. S15

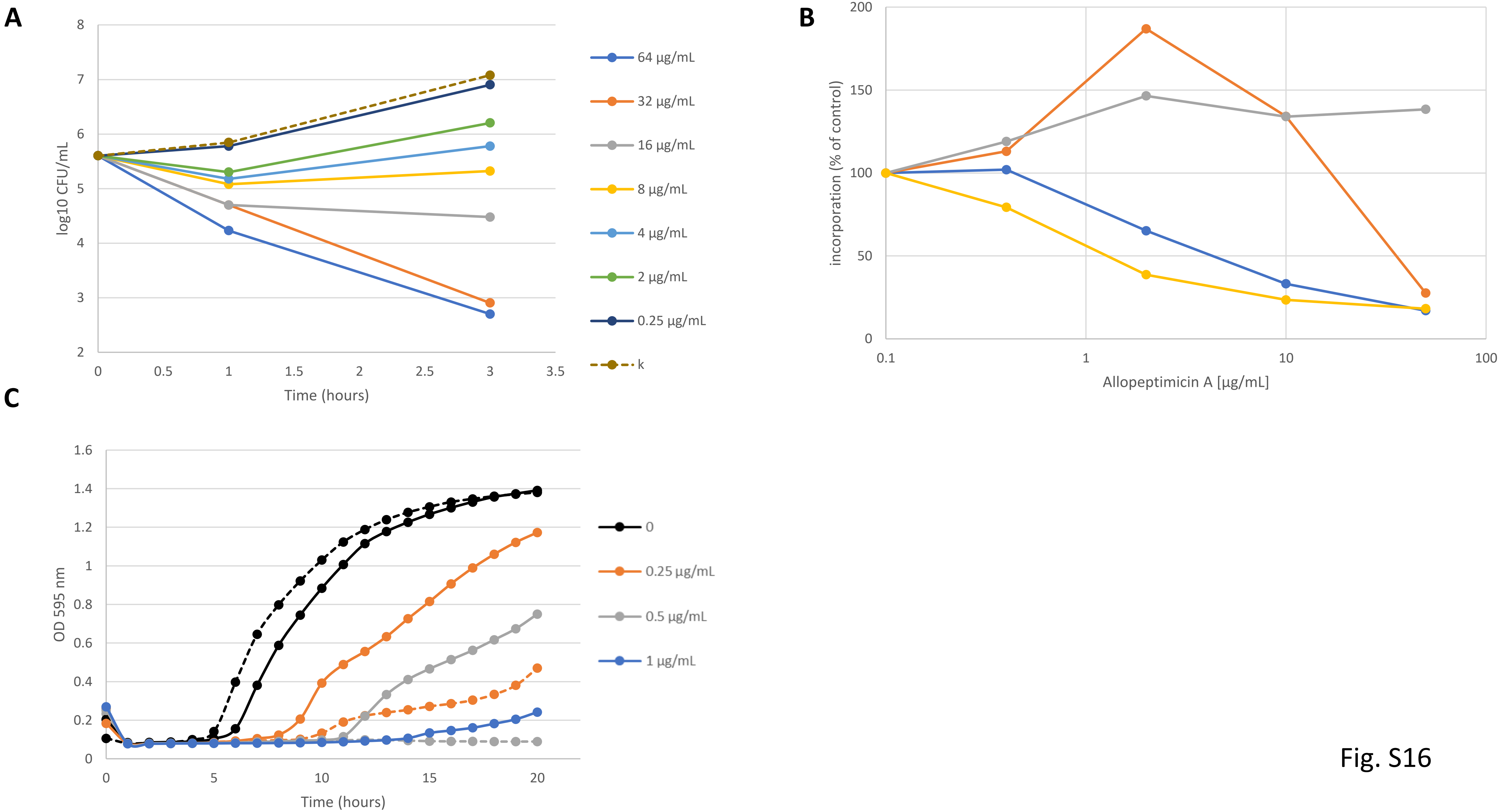

A
log10 CFU/mL
Time (hours)
64 µg/mL
32 µg/mL
16 µg/mL
8 µg/mL
4 µg/mL
2 µg/mL
0.25 µg/mL
k
B
incorporation (% of control)
Allopeptimicin A [µg/mL]
C
OD 595 nm
Time (hours)
0
0.25 µg/mL
0.5 µg/mL
1 µg/mL


Fig. S16

A

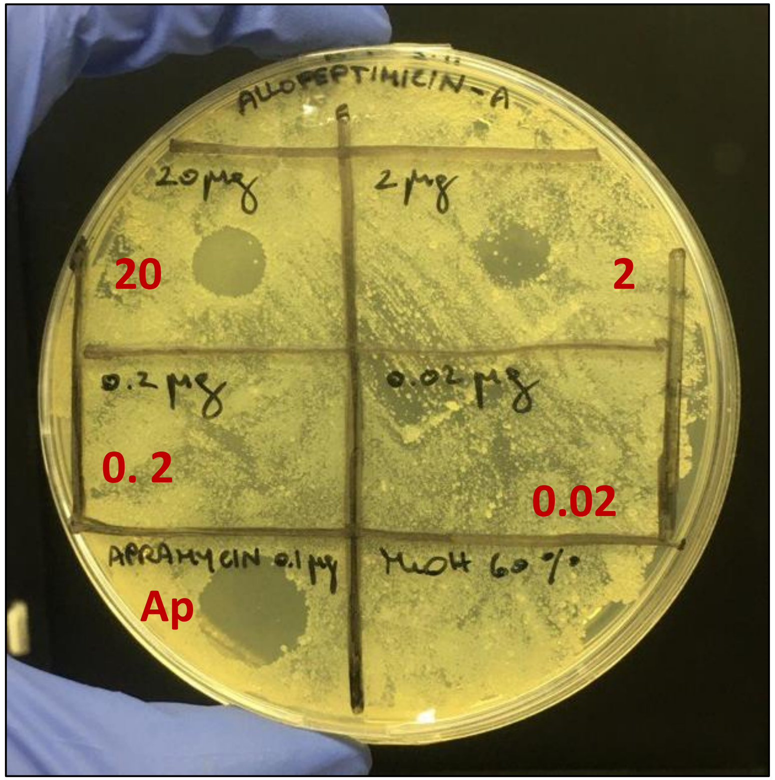
ALLOPEPTIMICIN-A
20 µg
2 µg
0.2 µg
0.02 µg
APRAMYCIN 0.1 µg
MeOH 60%
20
2
0. 2
0.02
Ap

B

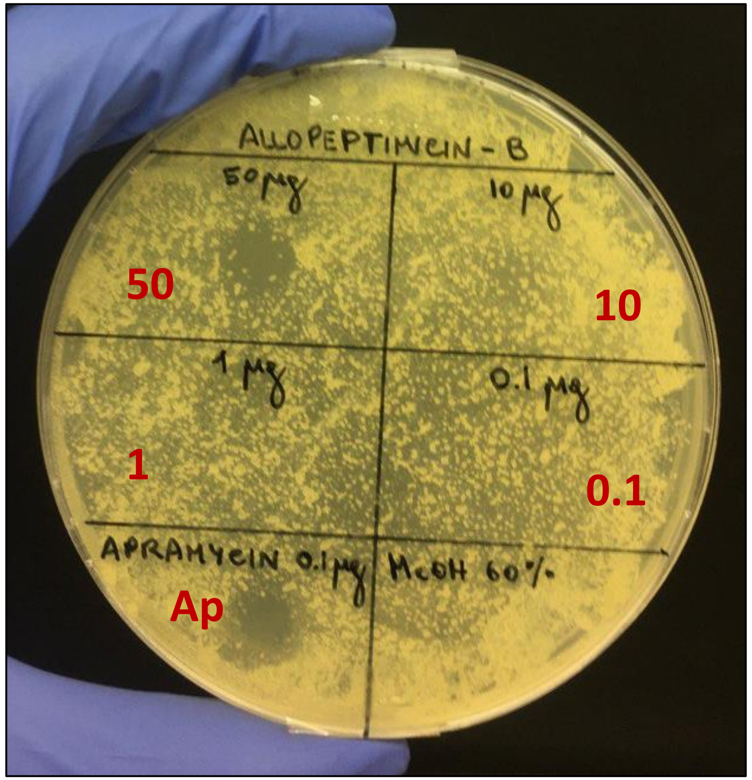
ALLOPEPTIMICIN-B
50 µg
10 µg
1 µg
0.1 µg
APRAMYCIN 0.1 µg
MeOH 60%
50
10
1
0.1
Ap

Fig. S17
